# Nonlinear ridge regression improves cell-type-specific differential expression analysis

**DOI:** 10.1101/2020.06.18.158758

**Authors:** Fumihiko Takeuchi, Norihiro Kato

## Abstract

**Background:** Epigenome-wide association studies (EWAS) and differential gene expression analyses are generally performed on tissue samples, which consist of multiple cell types. Cell-type-specific effects of a trait, such as disease, on the omics expression are of interest but difficult or costly to measure experimentally. By measuring omics data for the bulk tissue, cell type composition of a sample can be inferred statistically. Subsequently, cell-type-specific effects are estimated by linear regression that includes terms representing the interaction between the cell type proportions and the trait. This approach involves two issues, scaling and multicollinearity.

**Results:** First, although cell composition is analyzed in linear scale, differential methylation/expression is analyzed suitably in the logit/log scale. To simultaneously analyze two scales, we applied nonlinear regression. Second, we show that the interaction terms are highly collinear, which is obstructive to ordinary regression. To cope with the multicollinearity, we applied ridge regularization. In simulated data, nonlinear ridge regression attained well-balanced sensitivity, specificity and precision. Marginal model attained the lowest precision and highest sensitivity and was the only algorithm to detect weak signal in real data.

**Conclusion:** Nonlinear ridge regression performed cell-type-specific association test on bulk omics data with well-balanced performance. The omicwas package for R implements nonlinear ridge regression for cell-type-specific EWAS, differential gene expression and QTL analyses. The software is freely available from https://github.com/fumi-github/omicwas

## Background

Epigenome-wide association studies (EWAS) and differential gene expression analyses elucidate the association of disease traits (or conditions) with the level of omics expression, namely DNA methylation and gene expression. Thus far, tissue samples, which consist of heterogeneous cell types, have mainly been examined, because cell sorting is not feasible in most tissues and single-cell assay is still expensive. Nevertheless, the cell type composition of a sample can be quantified statistically by comparing omics measurement of the target sample with reference data obtained from sorted or single cells [1,2]. By utilizing the composition, the disease association specific to a cell type was statistically inferred for gene expression [3-10] and DNA methylation [11-14].

For the imputation of cell type composition, omics markers are usually analyzed in the original linear scale, which measures the proportion of mRNA molecules from a specific gene or the proportion of methylated cytosine molecules among all cytosines at a specific CpG site [15]. The proportion can differ between cell types, and the weighted average of cell-type-specific proportions becomes the proportion in a bulk tissue sample. Using the fact that the weight equals the cell type composition, the cell type composition of a sample is imputed. In contrast, gene expression analyses are performed in the log-transformed scale because the signal and noise are normally distributed after log-transformation [16]. In DNA methylation analysis, the logit-transformed scale, which is called the M-value, is statistically valid [17], although the linear scale could yield comparable performance under large sample size [18]. Consequently, the optimal scales for analyzing differential gene expression or methylation can differ from the optimal scale for analyzing cell type composition.

Aiming to perform cell-type-specific EWAS or differential gene expression analyses by using unsorted tissue samples, we study two issues that have been overlooked. Whereas previous studies were performed in linear scale, we develop a nonlinear regression, which simultaneously analyzes cell type composition in linear scale and differential expression/methylation in log/logit scale. The second issue is multicollinearity. Cell-type-specific effects of a trait, such as disease, on omics expression are usually estimated by linear regression that includes terms representing the interaction between the cell type proportions and the trait. We show that the interaction terms can mutually be highly correlated, which obstructs ordinary regression. To cope with the multicollinearity, we implement ridge regularization. Our methods and previous ones are compared in simulated and real data.

## Results

### Multicollinearity of interaction terms

Typically, cell-type-specific effects of a trait on omics marker expression is analyzed by the linear regression in equation (2). For each omics marker, the goal is to estimate *β*_*h,k*_, the effect of trait *k* on the expression level in cell type *h*. This is estimated based on the relation between the bulk expression level *Y*_*i*_ of sample *i* and the regressor *W*_*h,i*_*X*_*i,k*_, which is an interaction term defined as the product of the cell type proportion *W*_*h,i*_ and the trait value *X*_*i,k*_ of the sample. We assume that *Y*_*i*_, *W*_*h,i*_ and *X*_*i,k*_ are given as input data.

The variable *W*_*h,i*_ for cell type composition cannot be mean-centered for our purpose. If *W*_*h,i*_ were centered, we would obtain, instead of *β*_*h,k*_, the deviation of *β*_*h,k*_ from the average across cell types. In general, interaction terms involving uncentered variables can become collinear [19]. We first survey the extent of multicollinearity in real data for cell-type-specific association.

In peripheral blood leukocyte data from a rheumatoid arthritis study (GSE42861), the proportion of cell types ranged from 0.59 for neutrophils to 0.01 for eosinophils (Table 1A). The proportion of neutrophils was negatively correlated with the proportion of other cell types (apart from monocytes) with correlation coefficient of –0.68 to –0.46, whereas the correlation was weaker for other pairs (Table 1B). Rheumatoid arthritis status was modestly correlated with proportions of cell types. The product of the disease status *X*_*k*_, centered to have zero mean, and the proportion of a cell type becomes an interaction term. The correlation coefficients between the interaction terms were mostly >0.8, apart from eosinophils (Table 1C). The coefficient of variation (CV), which is the ratio of standard deviation to mean, of the proportion was low for all cell types apart from eosinophils (Table 1A). The interaction terms for low-CV cell types were strongly correlated with *X*_*k*_, which in turn caused strong correlation between the relevant interaction terms.

**Table 1A.** Blood cell type proportion in rheumatoid arthritis dataset

| Cell type | Neu | CD4<sup>+</sup>T | CD8<sup>+</sup>T | NK | Mono | Bcells | Eos |
| --- | --- | --- | --- | --- | --- | --- | --- |
| Mean | 0.59 | 0.10 | 0.08 | 0.08 | 0.07 | 0.07 | 0.01 |
| SD | 0.11 | 0.06 | 0.05 | 0.04 | 0.02 | 0.03 | 0.02 |
| CV | 0.2 | 0.6 | 0.6 | 0.5 | 0.3 | 0.4 | 2.7 |

SD, standard deviation; CV, coefficient of variation

**Table 1B.** Correlation between blood cell type proportion and rheumatoid arthritis (X_k_)

| $r$ | Neu | CD4<sup>+</sup>T | CD8<sup>+</sup>T | NK | Mono | Bcells | Eos | $X_k$=Disease |
| --- | --- | --- | --- | --- | --- | --- | --- | --- |
| **Neu** | 1 | -0.68 | -0.60 | -0.46 | -0.06 | -0.49 | -0.48 | 0.44 |
| **CD4<sup>+</sup>T** | -0.68 | 1 | 0.14 | 0.05 | -0.17 | 0.38 | 0.26 | -0.33 |
| **CD8<sup>+</sup>T** | -0.60 | 0.14 | 1 | 0.08 | -0.05 | 0.19 | 0.13 | -0.27 |
| **NK** | -0.46 | 0.05 | 0.08 | 1 | -0.04 | 0.01 | 0.11 | -0.27 |
| **Mono** | -0.06 | -0.17 | -0.05 | -0.04 | 1 | -0.17 | 0.05 | 0.10 |
| **Bcells** | -0.49 | 0.38 | 0.19 | 0.01 | -0.17 | 1 | 0.11 | -0.22 |
| **Eos** | -0.48 | 0.26 | 0.13 | 0.11 | 0.05 | 0.11 | 1 | -0.10 |

**Table 1C.** Correlation between interaction terms

| $r$ | Neu*$X_k$ | CD4<sup>+</sup>T*$X_k$ | CD8<sup>+</sup>T*$X_k$ | NK*$X_k$ | Mono*$X_k$ | Bcells*$X_k$ | Eos*$X_k$ |
| --- | --- | --- | --- | --- | --- | --- | --- |
| **Neu\*$X_k$** | 1 | 0.83 | 0.80 | 0.85 | 0.93 | 0.90 | 0.27 |
| **CD4<sup>+</sup>T\*$X_k$** | 0.83 | 1 | 0.78 | 0.78 | 0.83 | 0.88 | 0.42 |
| **CD8<sup>+</sup>T\*$X_k$** | 0.80 | 0.78 | 1 | 0.77 | 0.82 | 0.83 | 0.35 |
| **NK\*$X_k$** | 0.85 | 0.78 | 0.77 | 1 | 0.85 | 0.83 | 0.35 |
| **Mono\*$X_k$** | 0.93 | 0.83 | 0.82 | 0.85 | 1 | 0.88 | 0.35 |
| **Bcells\*$X_k$** | 0.90 | 0.88 | 0.83 | 0.83 | 0.88 | 1 | 0.36 |
| **Eos\*$X_k$** | 0.27 | 0.42 | 0.35 | 0.35 | 0.35 | 0.36 | 1 |

Neu, neutrophils; Mono, monocytes; Eos, eosinophils.

The situation was the same for the interaction with age in GTEx data. The granulocytes (which include neutrophils and eosinophils) were the most abundant (Table 2A). The proportion of granulocytes was negatively correlated with other cell types (apart from monocytes) with correlation coefficient of –0.89 to –0.41, and the correlation between other pairs was generally weaker (Table 2B). Age was modestly correlated with proportions of cell types. In this dataset, the CV of the proportion was low in all cell types (Table 2A), which caused strong mutual correlation between interaction terms (Table 2C).

**Table 2A.** Blood cell type proportion in GTEx dataset

| Cell type | Gran | CD4<sup>+</sup>T | CD8<sup>+</sup>T | Mono | NK | Bcells |
| --- | --- | --- | --- | --- | --- | --- |
| Mean | 0.53 | 0.22 | 0.10 | 0.07 | 0.05 | 0.03 |
| SD | 0.037 | 0.020 | 0.013 | 0.004 | 0.012 | 0.003 |
| CV | 0.1 | 0.1 | 0.1 | 0.1 | 0.2 | 0.1 |

SD, standard deviation; CV, coefficient of variation

**Table 2B.** Correlation between blood cell type proportion and age (X_k_)

| $r$ | Gran | CD4<sup>+</sup>T | CD8<sup>+</sup>T | Mono | NK | Bcells | $X_k$=Age |
| --- | --- | --- | --- | --- | --- | --- | --- |
| **Gran** | 1 | -0.89 | -0.83 | 0.56 | -0.76 | -0.41 | -0.23 |
| **CD4<sup>+</sup>T** | -0.89 | 1 | 0.59 | -0.64 | 0.50 | 0.51 | 0.14 |
| **CD8<sup>+</sup>T** | -0.83 | 0.59 | 1 | -0.40 | 0.59 | 0.15 | 0.15 |
| **Mono** | 0.56 | -0.64 | -0.40 | 1 | -0.44 | -0.42 | 0.02 |
| **NK** | -0.76 | 0.50 | 0.59 | -0.44 | 1 | 0.13 | 0.31 |
| **Bcells** | -0.41 | 0.51 | 0.15 | -0.42 | 0.13 | 1 | -0.03 |

**Table 2C.** Correlation between interaction terms

| $r$ | Gran*$X_k$ | CD4<sup>+</sup>T*$X_k$ | CD8<sup>+</sup>T*$X_k$ | Mono*$X_k$ | NK*$X_k$ | Bcells*$X_k$ |
| --- | --- | --- | --- | --- | --- | --- |
| **Gran\*$X_k$** | 1 | 0.99 | 0.98 | 1.00 | 0.96 | 0.99 |
| **CD4<sup>+</sup>T\*$X_k$** | 0.99 | 1 | 1.00 | 0.99 | 0.98 | 1.00 |
| **CD8<sup>+</sup>T\*$X_k$** | 0.98 | 1.00 | 1 | 0.99 | 0.98 | 0.99 |
| **Mono\*$X_k$** | 1.00 | 0.99 | 0.99 | 1 | 0.96 | 0.99 |
| **NK\*$X_k$** | 0.96 | 0.98 | 0.98 | 0.96 | 1 | 0.97 |
| **Bcells\*$X_k$** | 0.99 | 1.00 | 0.99 | 0.99 | 0.97 | 1 |

Gra, granulocytes; Mono, monocytes.

In the above empirical data, multicollinearity between interaction terms seemed to arise not due to the correlation between cell type proportions or *X*_*k*_, but due to the low CV in the cell type proportions. Subsequently, this property was derived mathematically. As we derived in equation (19), the correlation between interaction terms *W*_*h*_*X*_*k*_ and *W*_*h*′_ *X*_*k*_ approaches one when CV[*W*_*h*_] and CV[*W*_*h*′_] are low, irrespective of Cor[*W*_*h*_*M W*_*h*′_] (Fig. 1). The CV was 0.2 to 0.6 (apart from eosinophils) in the rheumatoid arthritis dataset and 0.1 to 0.2 in the GTEx dataset. We looked up datasets of several ethnicities and found the CV to be ≤0.6 in majority of blood cell types (Additional file 1: Table S1). Thus, multicollinearity can be a common problem for cell-type-specific association analyses. Biologically, in tissues where cell type composition is tightly controlled, the CV of cell type proportion becomes low and the multicollinearity is exacerbated.

**Figure 1.**
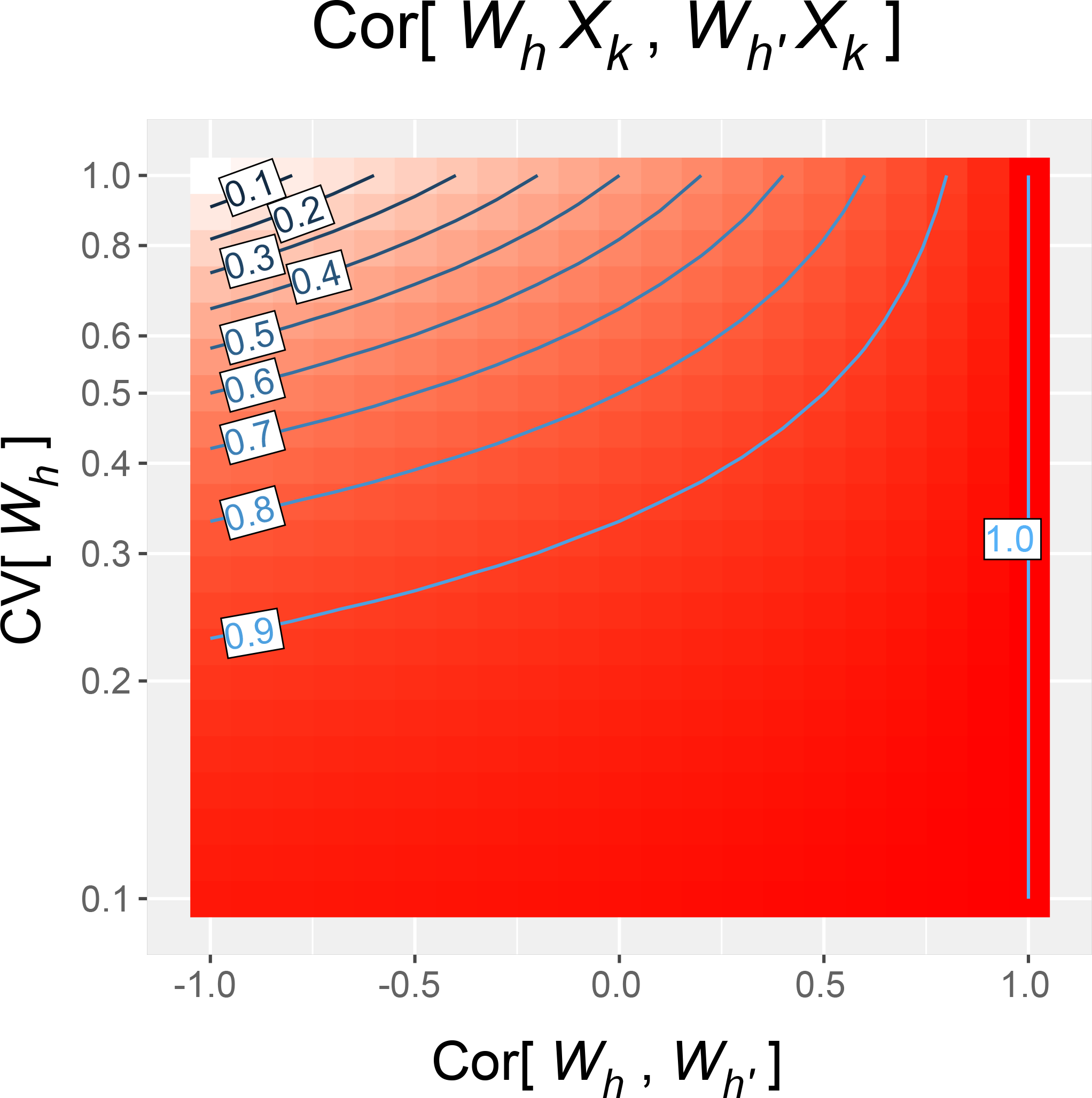
Contour plot of the correlation coefficient between interaction terms *W*_*h*_*X*_*k*_ and *W*_*h*′_ *X*_*k*_. *W*_*h*_ and *W*_*h*′_ represent proportions of cell types *h* and *h*^′^, and *X*_*k*_ represents the value of trait *k*. For this plot, we assume the coefficient of variation CV[*W*_*h*_] and CV[*W*_*h*′_] to be equal. As the CV decreases 0.6, 0.4 to 0.2, the correlation coefficient raises >0.5, >0.7 to >0.9, over most range of Cor[*W*_*h*_*M W*_*h*′_].

### Evaluation in simulated data

By using simulated data, we evaluated previous methods and new approaches of the omicwas package. In order to simultaneously analyze two scales, the linear scale for heterogeneous cell mixing and the log/logit scale for trait effects, we applied nonlinear regression (equations (4) and (5)). To cope with the multicollinearity of interaction terms, we applied ridge regularization (formula (10)).

Previous regression type methods are based either on the full model of linear regression (equation (2)) or the marginal model (equation (3)). The full model fits and tests cell-type-specific effects for all cell types simultaneously, and its variations include TOAST, csSAM.lm and CellDMC. The marginal model fits and tests cell-type-specific effect for one cell type at a time, and its variations include csSAM.monovariate and TCA. We also examined a hybrid of the two models (Marginal.Full005), which becomes positive when the models agree.

The simulation data was generated from real datasets of DNA methylation (658 samples; 451,725 CpG sites) and gene expression (389 samples; 14,038 genes). The original cell type composition was retained for all samples, and the case-control status was randomly assigned. Ninety-five percent of omics markers were set to be unassociated with disease status, 2.5% were up-regulated in cases at one cell type, and 2.5% were similarly down-regulated. The cell-type-specific effect-size was fixed in a simulation trial, either to methylation odds ratio (OR) of 1.3, 1.6 or 1.9 or to gene expression fold change of 1.7, 3.0 or 5.0. The significance level was set to P < 2.4 × 10^−7^ for DNA methylation and false discovery rate <5% for gene expression. In each simulation trial, the sensitivity, specificity and precision for detecting cell-type-specific association was calculated for each cell type. To compare algorithms, the performance measures for the same effect-size and cell type were averaged over the simulation trials.

Overall, in the simulation for DNA methylation the sensitivity (Fig. 2) was higher under large effect-size (bottom row of panels) and in abundant cell types (left columns of panels). The average specificity (Fig. 3) was high across the effect-size settings and across cell types, being >0.97 for the Marginal model, >0.98 for TCA and >0.999 for other algorithms. The precision (Fig. 4) was higher in abundant cell types within each effect-size setting. As effect-size increased, the precision decreased in neutrophils but increased in the other minor cell types. Excluding the cases where all algorithms lacked sensitivity (monocytes and B cells under methOR=1.3 and eosinophils), the average precision of omicwas.logit.ridge was >0.79 and was the highest in 13/16 of the cases.

**Figure 2.**
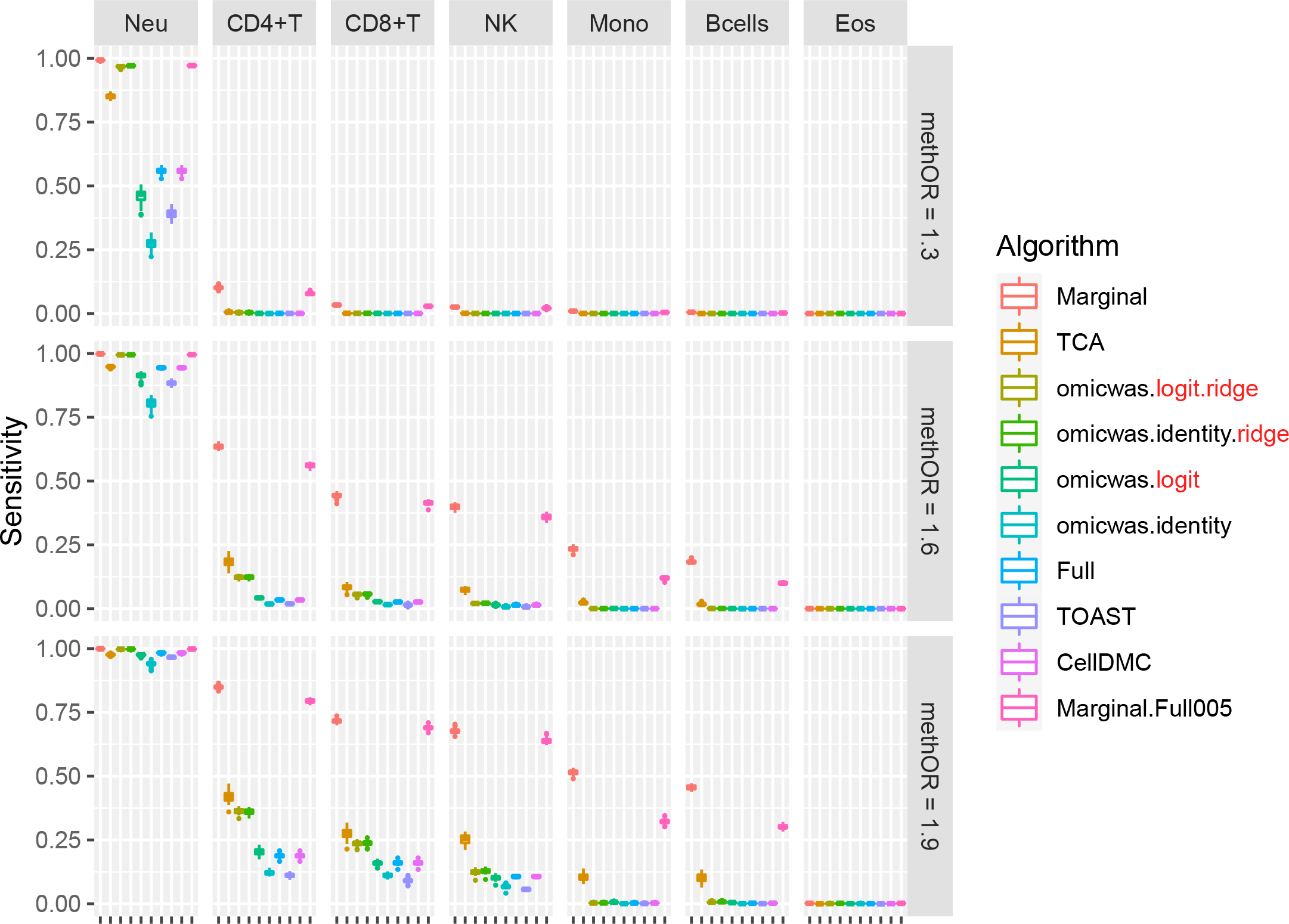
Sensitivity for detecting cell-type-specific association in simulated data for DNA methylation. Panels are aligned in rows according to the simulation settings with the methylation odds ratio of 1.3, 1.6 or 1.9. In each row, panels for different cell types are aligned in decreasing order of proportion. The vertical axis indicates sensitivity. In each panel, results from different algorithms are aligned horizontally in different colors. Results from 20 simulation trials are summarized in a box plot. The middle bar of the box plot indicates the median, and the lower and upper hinges correspond to the first and third quartiles. The whiskers extend to the value no further than 1.5 × inter-quartile range from the hinges. MethOR, methylation odds ratio; Neu, neutrophils; Mono, monocytes; Eos, eosinophils.

**Figure 3.**
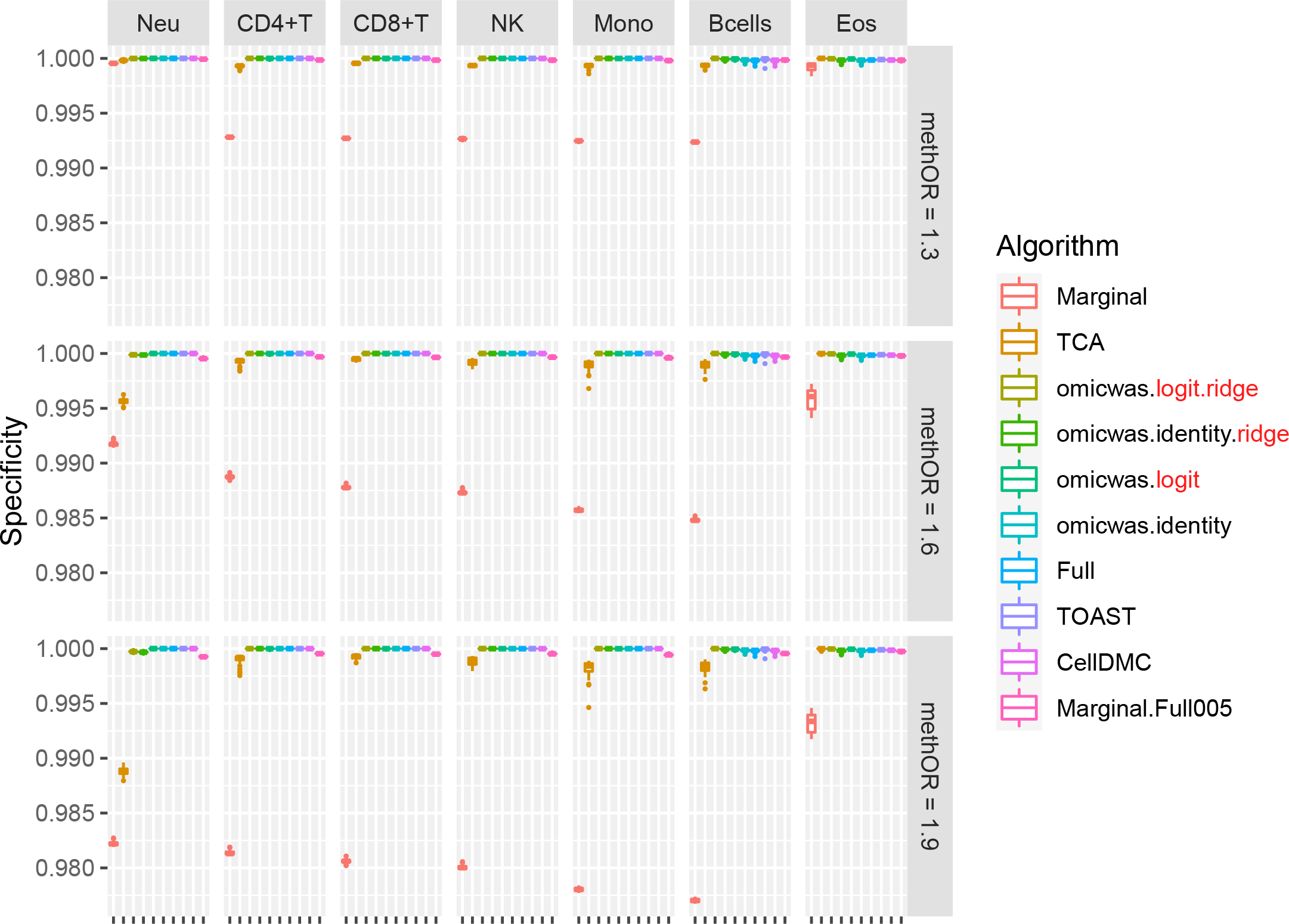
Specificity for detecting cell-type-specific association in simulated data for DNA methylation. The figure format is same as Fig. 3.

**Figure 4.**
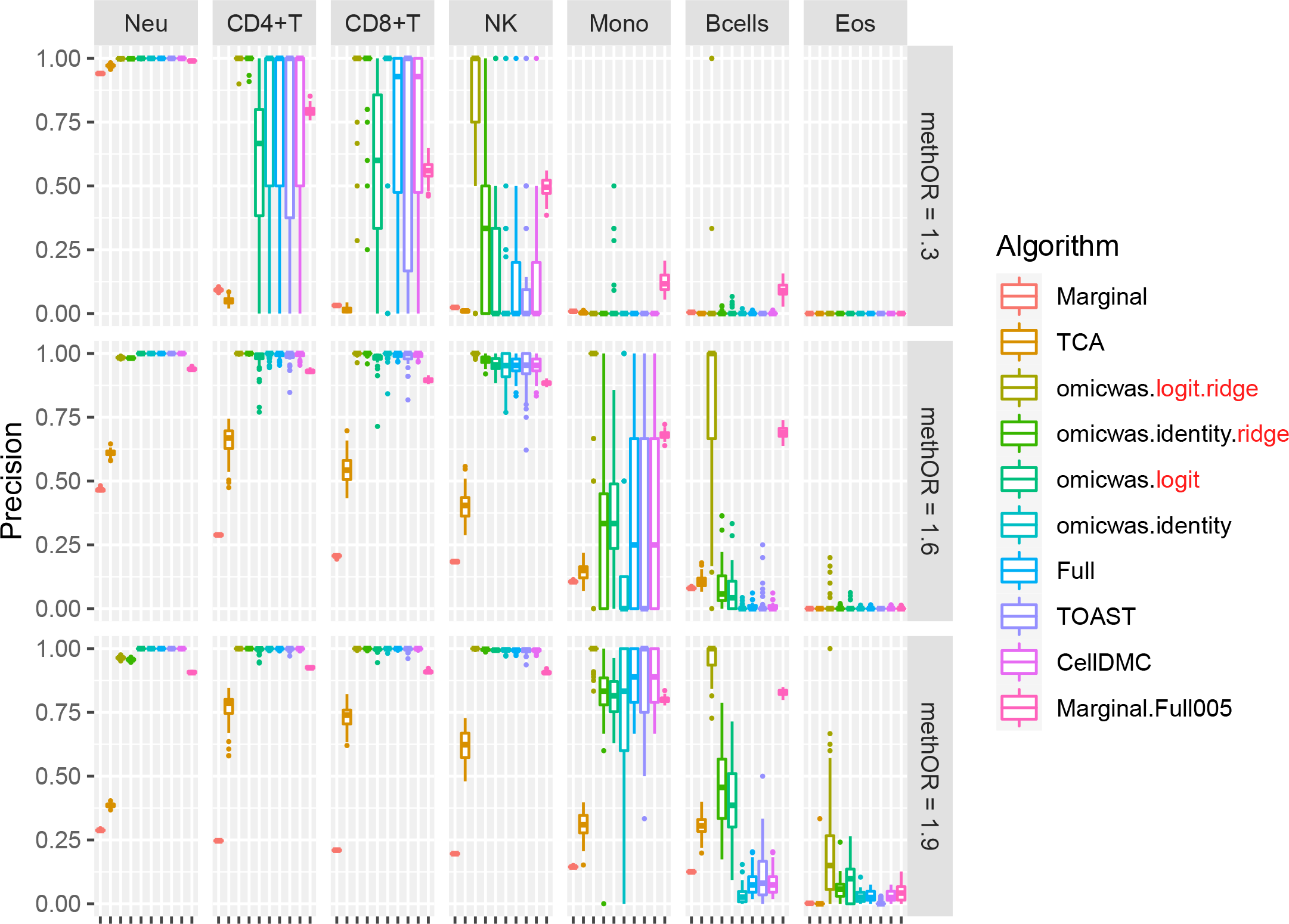
Precision (positive predictive value) for detecting cell-type-specific association in simulated data for DNA methylation. The figure format is same as Fig. 3.

There was trade-off between sensitivity and precision. Among the algorithms, the Marginal model attained the highest sensitivity and the lowest precision. TCA, which is a variation of the marginal model, had relatively high sensitivity and relatively low precision. Marginal.Full005 attained the second highest sensitivity and moderate precision. The ridge regressions (omicwas.logit.ridge and omicwas.identity.ridge) attained moderate sensitivity and high precision. The full models without ridge regularization (omicwas.logit, omicwas.identity, Full, TOAST and CellDMC) had the lowest sensitivity.

The overall tendency was similar in the simulation for gene expression. The sensitivity (Fig. 5) was higher under large effect-size and in abundant cell types. The average specificity (Fig. 6) was high across the effect-size settings and across cell types, being >0.96 for the marginal models (Marginal and csSAM.monovariate), >0.98 for the nonlinear and ridge regressions (omicwas.log.ridge, omicwas.identity.ridge, omicwas.log) and >0.996 for other algorithms. The precision (Fig. 7) was higher in abundant cell types within each effect-size setting. As effect-size increased, the precision decreased in granulocytes but increased in the other minor cell types. Excluding the full models (omicwas.identity, Full, TOAST, csSAM.lm) that lacked sensitivity, the algorithms that were frequently top in average precision were omicwas.identity.ridge (5 cases), omicwas.log (5 cases), omicwas.log.ridge (3 cases) and Marginal.Full005 (3 cases).

**Figure 5.**
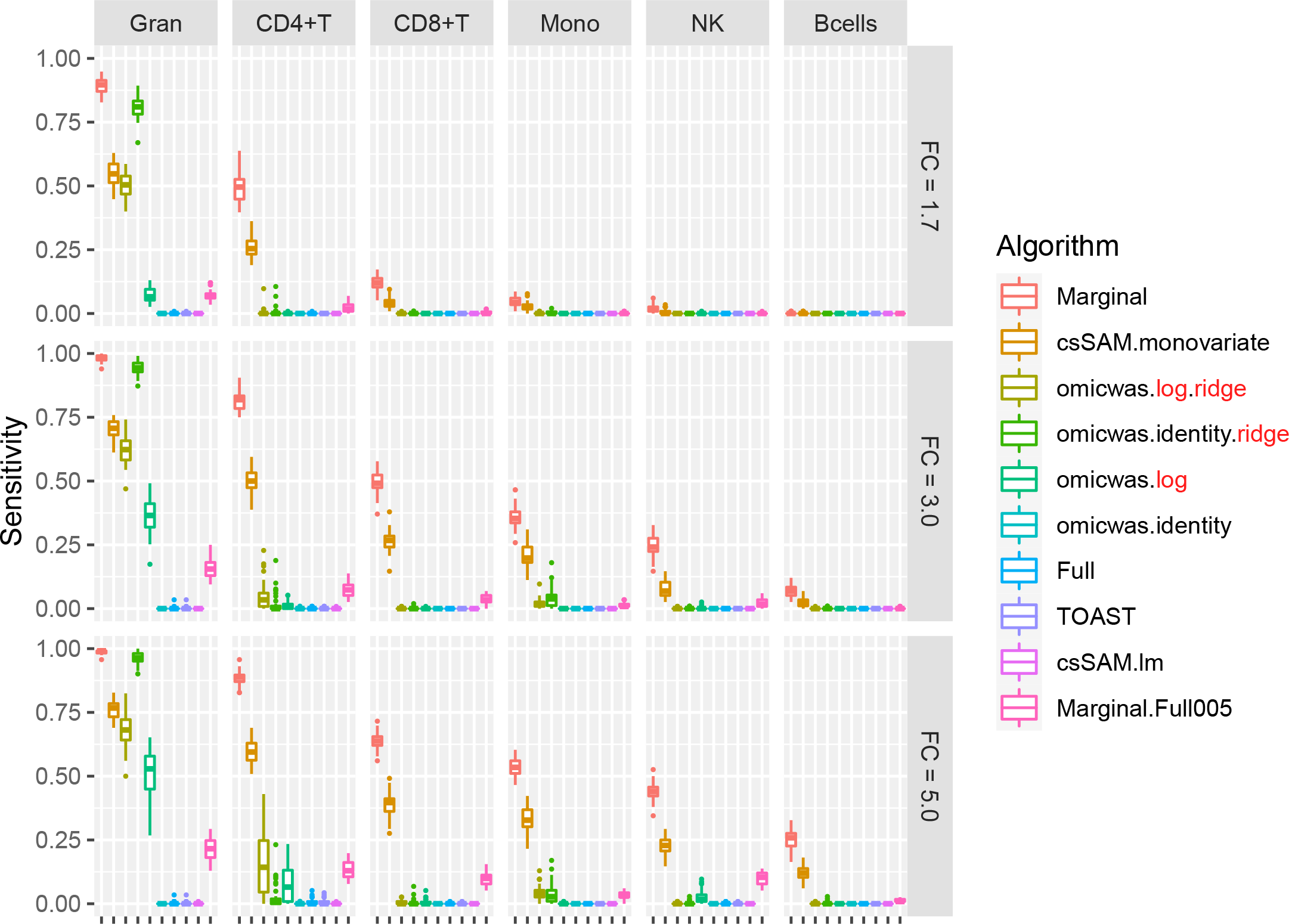
Sensitivity for detecting cell-type-specific association in simulated data for gene expression. Panels are aligned in rows according to the simulation settings with the gene expression fold change of 1.7, 3.0 or 5.0. In each row, panels for different cell types are aligned in decreasing order of proportion. The vertical axis indicates sensitivity. In each panel, results from different algorithms are aligned horizontally in different colors. Results from 50 simulation trials are summarized in a box plot. The middle bar of the box plot indicates the median, and the lower and upper hinges correspond to the first and third quartiles. The whiskers extend to the value no further than 1.5 × inter-quartile range from the hinges. FC, fold change; Gran, granulocytes; Mono, monocytes.

**Figure 6.**
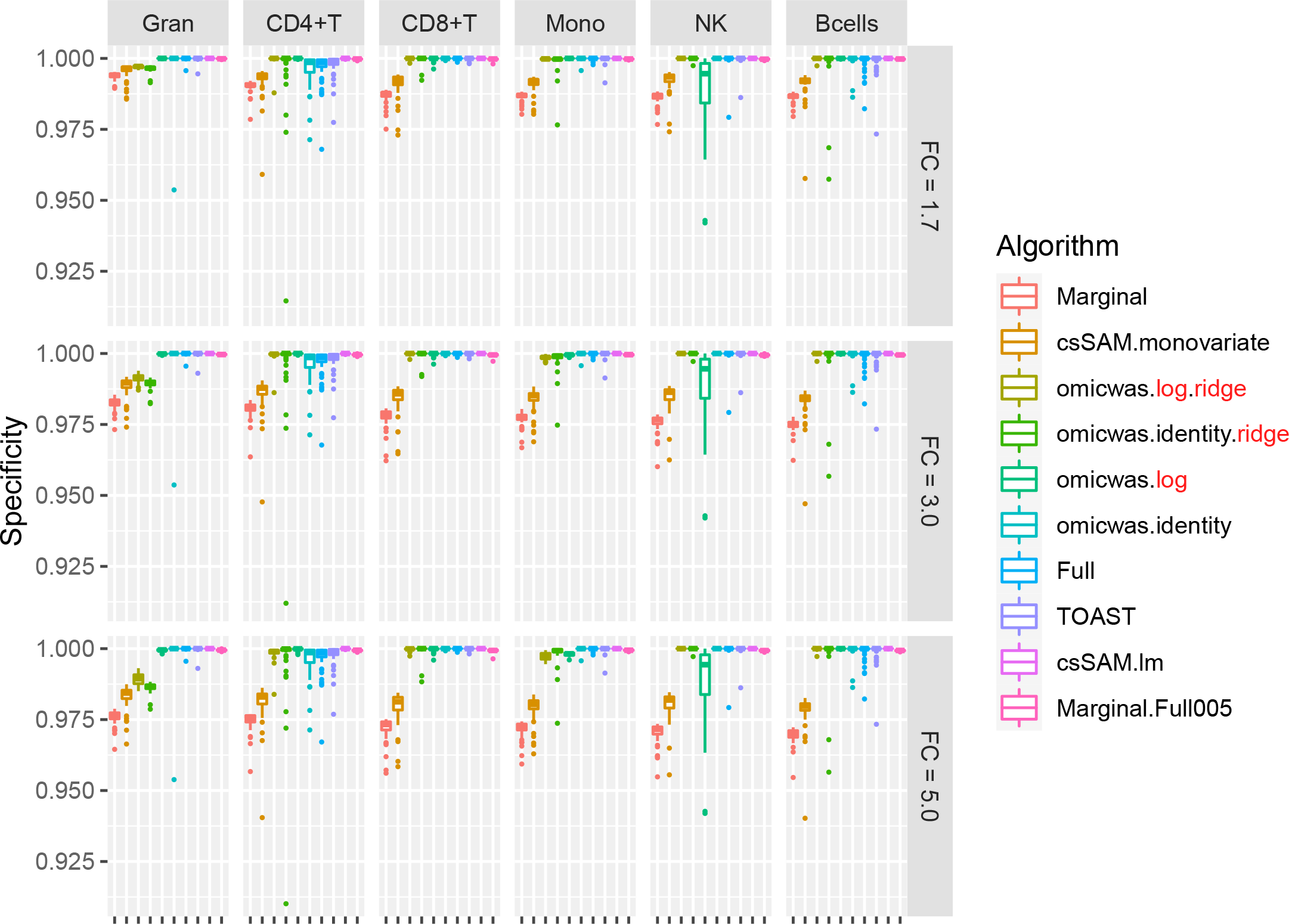
Specificity for detecting cell-type-specific association in simulated data for gene expression. The figure format is same as Fig. 5.

**Figure 7.**
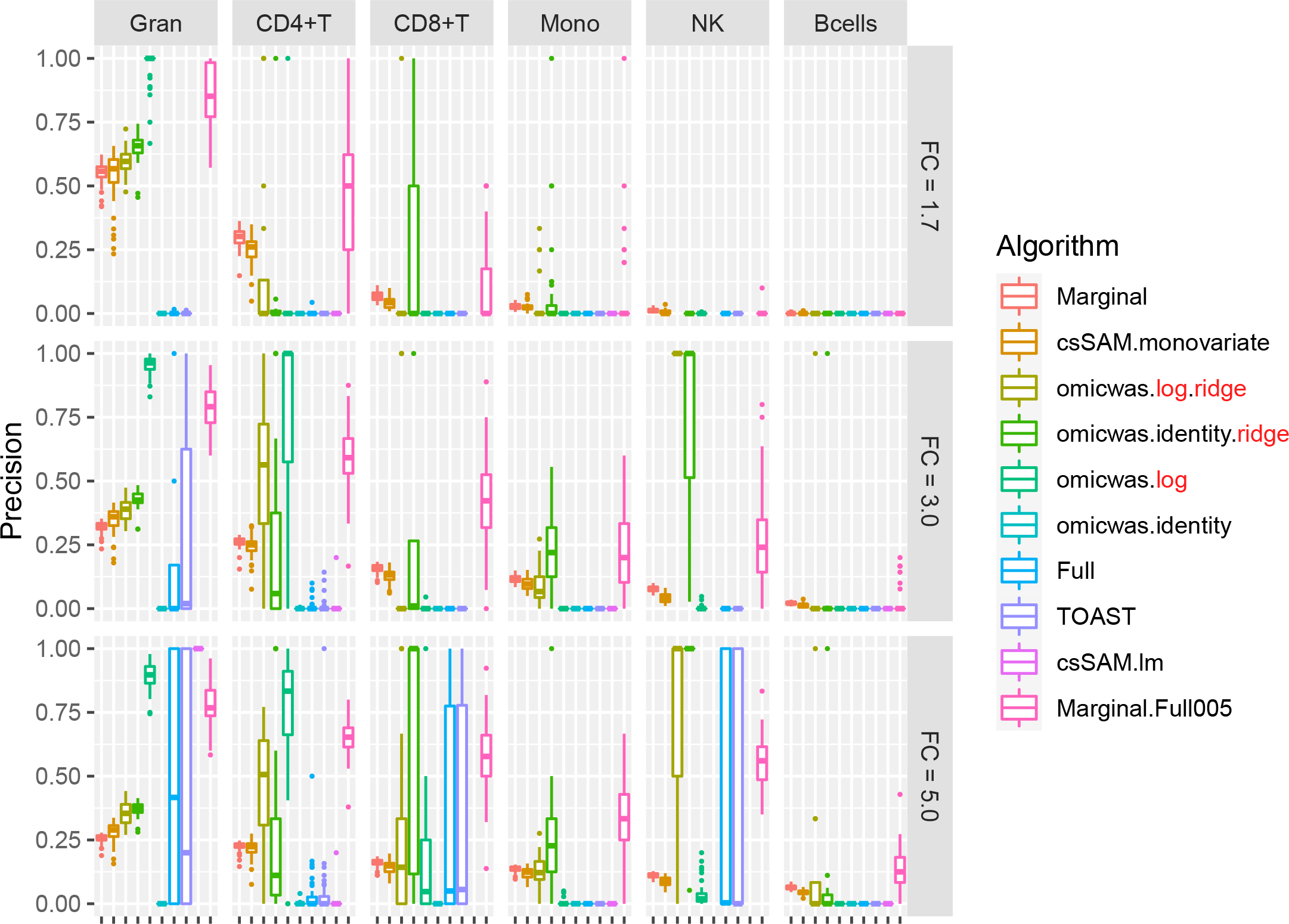
Precision (positive predictive value) for detecting cell-type-specific association in simulated data for gene expression. The figure format is same as Fig. 5.

There again was trade-off between sensitivity and precision. Among the algorithms, the marginal models attained the highest sensitivity but relatively low precision. The nonlinear and ridge regressions and Marginal.Full005 attained moderate sensitivity and highest precision. The full models had very low average sensitivity of <0.01.

For gene expression, we also simulated a scenario where cell-type-specific disease effect occurred in cell type “marker” genes (Additional files 2, 3, 4: Figs. S1, S2, S3). In other words, the expression level in the target cell type differed between cases and controls, and the expression level in other cell types was zero (in linear scale). Thus, non-target cell types did not introduce noise to bulk expression level. The Marginal model attained the highest average sensitivity of >0.93 but relatively low average precision of ∼0.16. As equal number of differentially expressed genes were generated in the six cell types, picking up signals for all such genes, including those not for the tested cell type, would result in precision of 1/6 = 0.16. The full models had low sensitivity. The nonlinear and ridge regressions and Marginal.Full005 attained moderate sensitivity and moderate precision. With regards to the frequency of being the top in average precision, the algorithms were ordered Marginal.Full005 (9 cases), omicwas.log.ridge (6 cases), omicwas.identity.ridge (2 cases) and omicwas.log (1 case), excluding the full models that lacked sensitivity.

### Cell-type-specific association with rheumatoid arthritis and age

The detection of cell-type-specific association in bulk tissue was evaluated by using physically sorted cells. In principle, sorted cells should serve as genuine verification, however, due to the relatively small sample size (94 or 203 for rheumatoid arthritis and 214 or 1202 for age) the available datasets were underpowered to generate a gold standard list of differentially expressed omics markers [20]. Instead, we generated a benchmark set of differentially expressed markers by imposing a relaxed significance level of P < 0.05; the set would be enriched for true differentially expressed markers yet also include unassociated markers. The benchmark set was cross-checked with the prediction by each algorithm; in the same manner as the simulation analysis, we assessed the sensitivity, specificity and precision.

The cell-type-specific association of DNA methylation with rheumatoid arthritis was predicted using bulk peripheral blood leukocyte data and was evaluated in sorted monocytes and B cells (Fig. 8). The input bulk methylation data was normalized by applying the logit-transformation for the Marginal.logit algorithm, which otherwise was the same as Marginal. Although the sensitivity was extremely low for all algorithms, it was positive in both cell types for Marginal (0.8–1.1 × 10^−4^), Marginal.logit (1.2–1.4 × 10^−4^), TCA (0.5–1.2 × 10^−4^) and omicwas.logit (0.5 × 10^−4^). The cell-type-specific association of DNA methylation with age was predicted using the same bulk dataset and was evaluated in sorted CD4^+^T cells and monocytes (Fig. 8). The Marginal and Marginal.logit models attained by far the highest sensitivity (both 0.15–0.27) in both cell types, and moderate precision (0.59–0.68 and 0.60–0.68 respectively).

**Figure 8.**
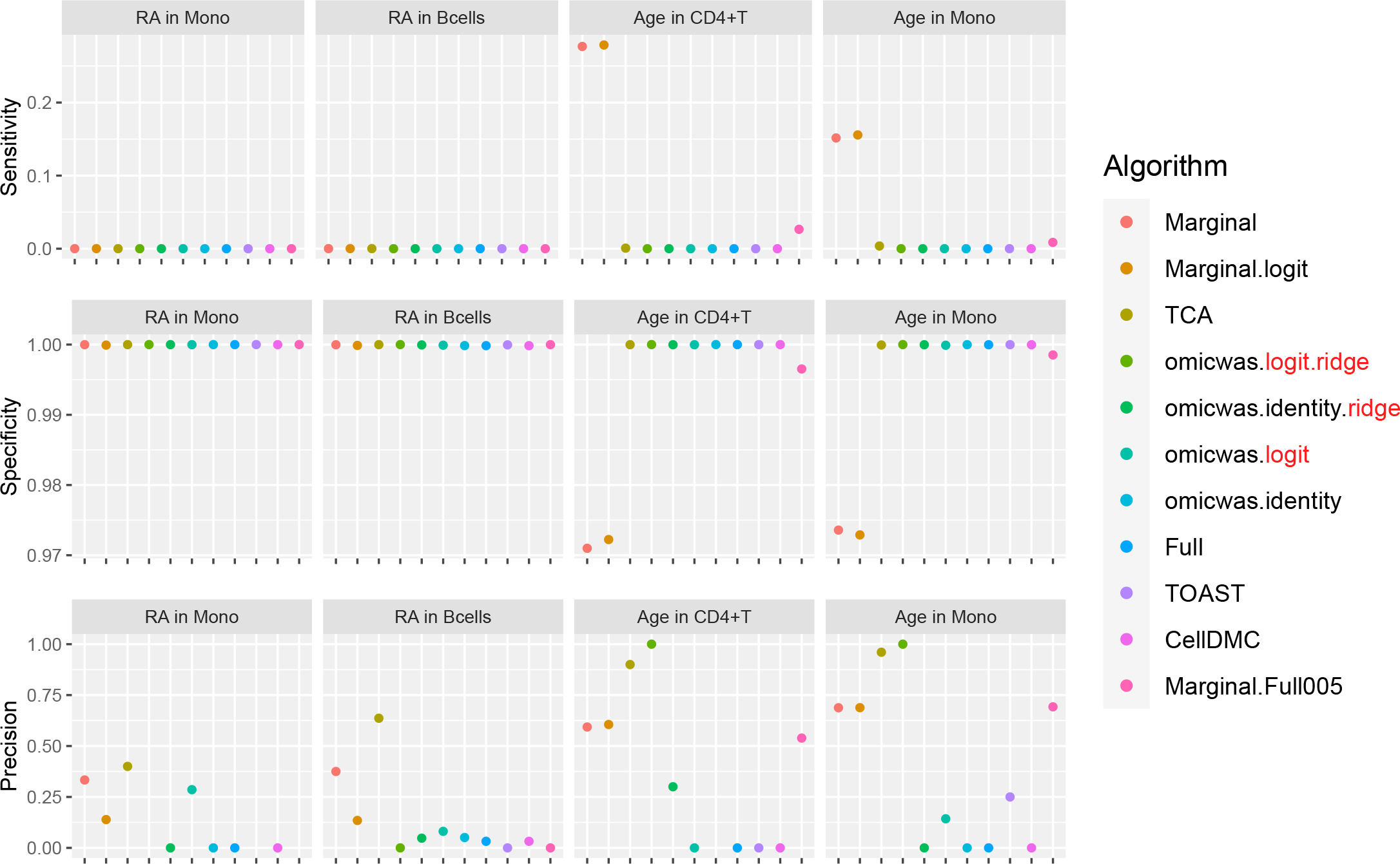
Performance of the predictions for cell-type-specific association of DNA methylation. For the association with rheumatoid arthritis in monocytes and B cells and the association with age in CD4^+^ T cells and monocytes, sensitivity (top), specificity (middle) and precision (bottom) are plotted. In each panel, results from different algorithms are aligned horizontally in different colors. Precision is not plotted when there were no positive CpG sites. RA, rheumatoid arthritis; Mono, monocytes.

The cell-type-specific association of gene expression with age was predicted using whole blood data and was evaluated in sorted CD4^+^ T cells and monocytes (Fig. 9). The input bulk gene expression data was normalized by applying the log-transformation for the Marginal.log algorithm, which otherwise was the same as Marginal. Although the sensitivity was low for all algorithms, it was positive in both cell types for Marginal (0.02–0.07), Marginal.log (0.07–0.11), omicwas.identity.ridge (0.01–0.22) and omicwas.log (0.03–0.05). The precision was modest for Marginal (0.06–0.31), Marginal.log (0.07–0.28), omicwas.identity.ridge (0.03–0.21) and omicwas.log (0.04–0.17). The dataset of sorted CD4^+^ T cells (214 samples) is smaller than the monocyte dataset (1202 samples) thus could be underpowered to pick enough true differentially expressed genes into the benchmark set.

**Figure 9.**
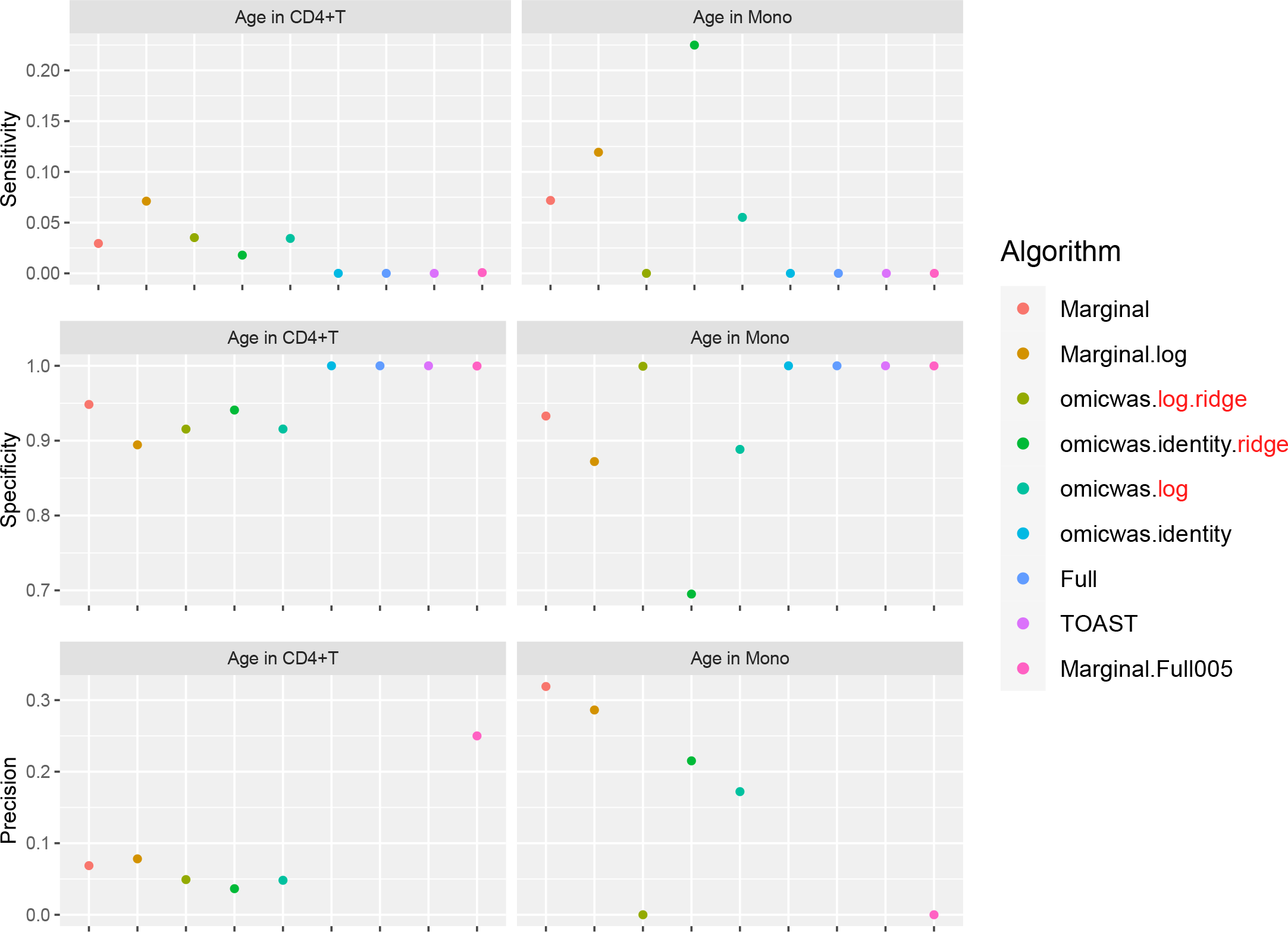
Performance of the predictions for cell-type-specific association of gene expression. For the association with age in CD4^+^ T cells and monocytes, sensitivity (top), specificity (middle) and precision (bottom) are plotted. In each panel, results from different algorithms are aligned horizontally in different colors. Precision is not plotted when there were no positive genes. Mono, monocytes.

For DNA methylation dataset GSE42861 and for GTEx gene expression dataset, the omicwas.logit.ridge and omicwas.log.ridge models of the omicwas package was computed in 8.1 and 0.7 hours respectively, using 8 cores of a 2.5 GHz Xeon CPU Linux server.

## Discussion

Aiming to elucidate cell-type-specific trait association in DNA methylation and gene expression, this article explored two aspects, multicollinearity and scale. We observed multicollinearity in real data and derived mathematically how it emerges. To cope with the multicollinearity, we applied ridge regularization. To properly handle multiple scales simultaneously, we applied nonlinear regression. Among the examined algorithms, nonlinear ridge regression attained moderate sensitivity and highest precision in simulated data. We also developed an algorithm that combines full and marginal models, which attained balanced sensitivity and precision in simulation. In real benchmark data, all algorithms performed poorly yet the marginal models tended to attain the highest sensitivity.

The statistical methods discussed in this article are applicable, in principle, to any tissue. For validation of the methods, we need datasets for bulk tissue as well as sorted cells, ideally of several hundred samples. Currently, the publicly available data is limited to peripheral blood. By no means, the rheumatoid arthritis EWAS datasets [21-23] or the datasets for age association of gene expression [24,25] are representative. Nevertheless, we think verification in real data is valuable.

By the performance in simulated and real data, we can roughly divide algorithms into three groups: full (and its variations), marginal (and its variations) and the third group that includes ridge regressions and the hybrid Marginal.Full005. In marginal models, we test one cell type at a time. If we knew in advance that one particular cell type is associated with the trait, which would be a rare situation, testing that cell type with the marginal model is the most simple and correct approach. However, when the test target cell type is not associated, but instead another cell type is associated, the marginal models can pick up false signals due to the collinearity between regressor variables. Indeed, marginal models attained highest sensitivity (Figs. 2, 5) and relatively low precision (Figs. 4, 7), which could lead to unstable performance. The full model fits and tests all cell types simultaneously, by which it adjusts for the effects of other cell types. Due to the simultaneous inclusion of collinear predictors, the sensitivity was low (Figs. 2, 5). The ridge regressions (omicwas.identity.ridge, omicwas.logit.ridge and omicwas.log.ridge) were in the middle between full and marginal models with regards to the sensitivity (Figs. 2, 5), while attaining the highest specificity (Figs. 4, 7). The hybrid Marginal.Full005 algorithm is intended to gain sensitivity by the marginal model while keeping precision by incorporating the full model. It attained moderate sensitivity (Figs. 2, 5) and moderate precision (Figs. 4, 7) in simulation. In real data, all algorithms performed poorly yet Marginal, Marginal.logit and Marginal.log tended to attain the highest sensitivity. With regards to the performance measures of all algorithms, the association of DNA methylation with age (Fig. 8) was roughly similar to the simulation setting of methylation OR = 1.6 for B cells (Figs. 2– 4), and the association of gene expression with age (Fig. 9) was roughly similar to the simulation setting of fold change = 1.7 for CD8^+^T cells (Figs. 5–7). For the respective simulation settings, the median coefficient of determination for the Marginal model was 0.020 and 0.007, indicating weak association.

A limitation of our simulation is that only one cell type was assumed to be associated with disease status at each marker. In reality, two or more cell types can be associated with disease under homogeneous or heterogeneous effect. In the physically sorted cells, the association of DNA methylation with rheumatoid arthritis tended to be consistent between monocytes and B cells; the association statistics across CpG sites were positively correlated with Spearman’s rank correlation coefficient of 0.20 (P-value < 2.2 × 10^−16^). Similarly, the association with age tended to be consistent between CD4^+^T cells and monocytes with correlation coefficient of 0.27 and 0.07 (P-value < 2.2 × 10^−16^), respectively, for DNA methylation and gene expression. The consistency suggests that multiple cell types tend to be associated under homogeneous effect. If the association is completely consistent, the effect-size is uniform across cell types. As there is no cell-type-specific effect, a simple regression by disease (or relevant trait), ignoring the cell type composition, becomes the appropriate modeling (formula (8)). Moreover, when cell type composition has low CV (as observed in Tables 1 and 2), the marginal model with normalized input (formula (9)) becomes almost identical to the simple regression. In other words, the marginal model can pick up signal in cases where effect-size is homogeneous across cell types. Correspondingly, in real data of DNA methylation Marginal.logit performed the best and was slightly better than Marginal (Fig. 8), and in gene expression Marginal.log performed mostly the best and was better than Marginal (Fig. 9).

## Conclusions

For cell-type-specific differential expression analysis by using unsorted tissue samples, we recommend trying the nonlinear ridge regression as a first choice because it balances sensitivity and precision. Although marginal models can be powerful when the tested cell type actually is the only one associated with the trait, caution is needed in its low precision. Under the idea of first scanning by the marginal model and then reanalyzing in full model, we developed the hybrid Marginal.Full005 algorithm, which attained balanced sensitivity and precision but was not corroborated in experimental data. Ridge regression is preferable compared to the full model without ridge regularization because ridge estimator of the effect-size has smaller mean squared error (equation (15)). The number of cell types associated with disease at each marker was restricted to one in our simulation but could be two or more with homogeneous or heterogeneous effect. If the effect-size is uniform across all cell types, a simple regression by disease status is suitable, which can be substituted with the marginal model that takes normalized input. We do not claim the ridge regression to substitute previous algorithms. Indeed, we think none of the current algorithms is superior to others in all aspects, indicating possibility for future improvement.

## Methods

### Linear regression

We begin by describing the linear regressions used in previous studies. Let the indexes be *h* for a cell type, *i* for a sample, *j* for an omics marker (CpG site or gene), *k* for a trait that has cell-type-specific effects on marker expression, and *l* for a trait that has a uniform effect across cell types. The input data is given in four matrices. The matrix *W*_*h,i*_ represents cell type composition. The matrices *X*_*i,k*_ and *C*_*i,l*_ represent the values of the traits that have cell-type-specific and uniform effects, respectively. We assume the two matrices are centered: Σ_*i*_ *X*_*i,k*_ = Σ_*i*_ *C*_*i,l*_ = 0. For example, *X*_*i,k*_ = 0.5 for disease cases and *X*_*i,k*_ = −0.5 for controls when the number of cases and controls are equal. The matrix *Y*_*i,j*_ represents the omics marker expression level in tissue samples.

The parameters we estimate are the cell-type-specific trait effect *β*_*h,jMk*_, tissue-uniform trait effect *γ*_*j,l*_, and basal marker level *α*_*h,j*_ in each cell type. For the remaining of the first five sections (up to “Multicollinearity of interaction terms”), we focus on one marker *j*, and omit the index for readability. For cell type *h*, the marker level of sample *i* is

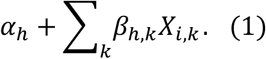

This is a representative value rather than a mean because we do not model a probability distribution for cell-type-specific expression. By averaging the value over cell types with weight *W*_*h,i*_, and combining with the tissue-uniform trait effects, we obtain the mean marker level in bulk tissue of sample *i*,

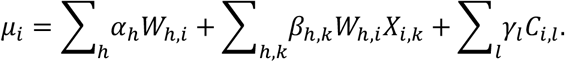

With regards to the statistical model, we assume the error of the marker level to be normally distributed with variance *σ*^2^, independently among samples, as

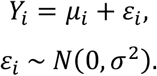

The statistical significance of all parameters is tested under the *full* model of linear regression,

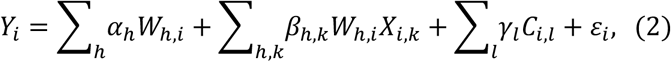

or its variations [5,9,13]. Alternatively, the cell-type-specific effects of traits can be fitted and tested for one cell type *h* at a time by the *marginal* model,

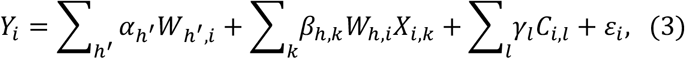

or its variations [7,8,10,11,14].

### Nonlinear regression

Aiming to simultaneously analyze cell type composition in linear scale and differential expression/methylation in log/logit scale, we develop a nonlinear regression model. The differential analyses are performed after applying normalizing transformation. The normalizing function is the natural logarithm *f* = log for gene expression, and *f* = logit for methylation (see Background). Conventional linear regression can be formulated by defining *f* as the identity function. We denote the inverse function of *f* by *g*; *g* = exp for gene expression, and *g* = logistic for methylation. Thus, *f* converts from the linear scale to the normalized scale, and *g* does the opposite.

The marker level in a specific cell type (formula (1)) is modeled in the normalized scale. The level is linearized by applying function *g*, then averaged over cell types with weight *W*_*h,i*_, and normalized by applying function *f*. Combined with the tissue-uniform trait effects, the mean normalized marker level in bulk tissue of sample *i* becomes

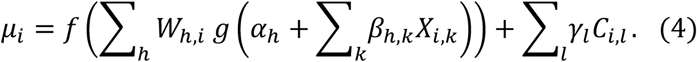

We assume the normalized marker level to have an error that is normally distributed with variance *σ*^2^, independently among samples, as

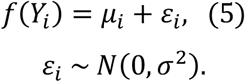

We obtain the ordinary least squares (OLS) estimator of the parameters by minimizing the residual sum of squares,

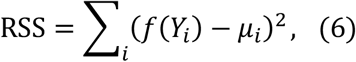

and then estimate the error variance as

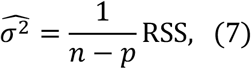

where *n* is the number of samples and *p* is the number of parameters [[26], section 6.3.1].

In the special case where the marker expression is homogeneous across cell types, the formulae become simple. Suppose that *α*_*h*_ regardless of cell type *h* equals *α* and that *β*_*h,k*_ equals *β*_*k*_. The regression formulae (4) and (5) of sample *i* reduces to

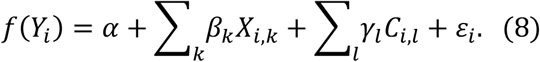

On the other hand, the marginal model for cell type *h* in formula (3) reduces to

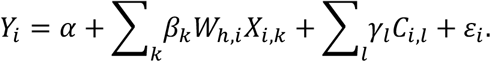

Moreover, when the CV for cell type composition is low, the cell type proportion *W*_*h,i*_ of sample *i* approximately equals the average 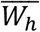 taken over samples. Thus, the formula reduces further to

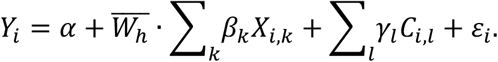

If we replace the input bulk expression level *Y*_*i*_ with the normalized value *f*(*Y*_*i*_), the model becomes

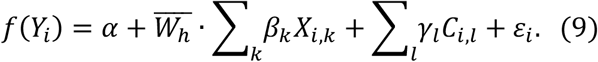

Under the special case of cell-type-homogeneous expression and low-CV cell type composition, formula (8) for nonlinear regression and formula (9) for the marginal model with normalized input become almost identical. The difference is the multiplication by constant 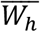, which does not change the test statistics for *β*_*k*_.

### Ridge regression

The parameters *β*_*h,k*_ for cell-type-specific effect cannot be estimated accurately by ordinary linear regression because the regressors *W*_*h,i*_*X*_*i,k*_ in equation (2) are highly correlated between cell types (see below). Multicollinearity also occurs to the nonlinear case in formula (4) because of local linearity. To cope with the multicollinearity, we apply ridge regression with a regularization parameter *λ* ≥ 0, and obtain the ridge estimator of the parameters that minimizes

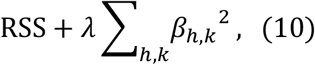

where the second term penalizes *β*_*h,k*_ for taking large absolute values. The ridge estimator 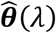 is asymptotically normally distributed (see Additional file 5: Supplementary note) with

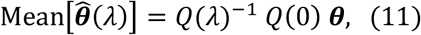

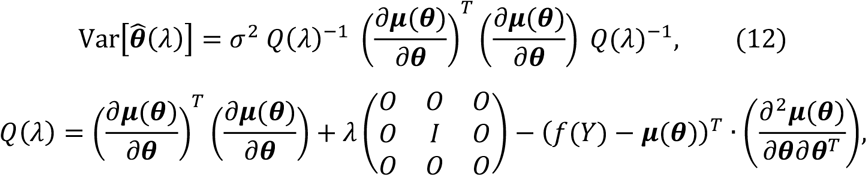

where ***μ*** is the vector form of *μ*_*i*_, ***θ*** is the vector form of the parameters *α*_*h*_, *β*_*h,k*_ and *γ*_*l*_ combined, (*∂****μ***/ *∂****θ***1 is the Jacobian matrix, (*∂*^2^***μ***/*∂****θ****∂****θ***^*T*^) is the array of Hessian matrices for *μ*_*i*_ taken over samples, and superscript *T* indicates matrix transposition. The dot product of (*f*(*Y*) − ***μ***(***θ***))^*T*^ and the array of Hessians is taken by multiplying for each sample and then summing up over samples. The matrix after *λ* has one only in the diagonal corresponding to *β*_*h,k*_. The assigned value ***θ*** is the true parameter value. By taking the expectation of *Q*, we obtain a rougher approximation [27] as

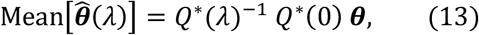

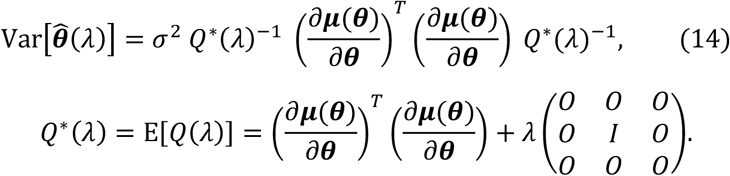

The matrices *Q* and *Q*^∗^ are the observed and expected Fisher matrices multiplied by *σ*^2^ and adapted to ridge regression, respectively.

Since our objective is to predict the cell-type-specific trait effects, we choose the regularization parameter *λ* that can minimize the mean squared error (MSE) of *β*_*h,k*_. Our methodology is based on [28]. To simplify the explanation, we assume the Jacobian matrices (*∂****μ***(***θ***)/ *∂****α***), (*∂****μ***(***θ***)/ *∂****β***) and (*∂****μ***(***θ***)/ *∂****γ***) to be mutually orthogonal, where ***α, β*** and ***γ*** are the vector forms of *α*_*h*_, *β*_*h,k*_ and *γ*_*l*_, respectively. Then, from formulae (13) and (14), the ridge estimator 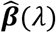 is asymptotically normally distributed with

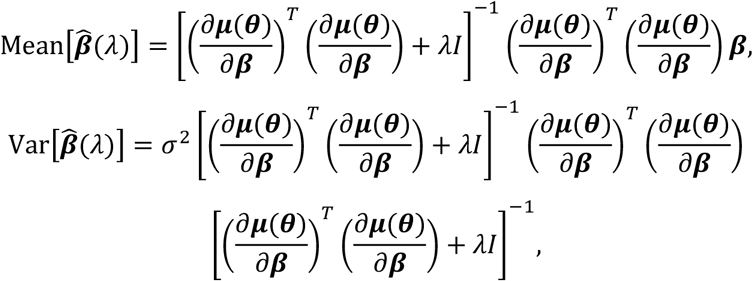

where the assigned values ***θ*** and ***β*** are the true parameter values. We apply singular value decomposition

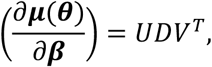

where *U* and *V* are orthogonal matrices, the columns of *V* are ***ν***_1_, …, ***ν***_*M*_, and the diagonals of diagonal matrix *D* are sorted *d*_1_ ≥ … ≥ *d*_*M*_ ≥ 0. The bias, variance and MSE of the ridge estimator are decomposed as

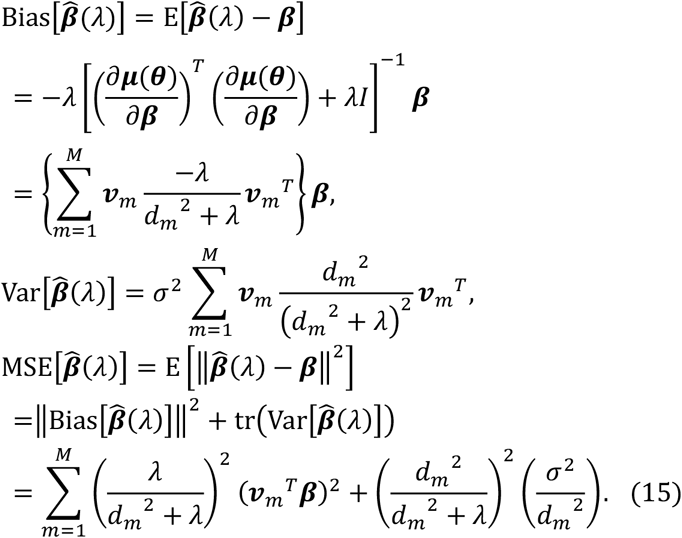

For each *m* in the summation of (13), the minimum of the summand is attained at *λ*_*m*_ = *σ*^2^/ (***ν***_*m*_^*T*^***β***)^2^. To minimize MSE, we need to find some “average” of the optimal *λ*_*m*_ over the range of *m*. Hoerl et al. [29] proposed to use the harmonic mean *λ* = *Mσ*^2^/ ‖***β***‖^2^. However, if an OLS estimator 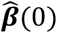 is actually plugged into ‖***β***‖^2^, the denominator is biased upwards, and the computed mean is biased downwards. Indeed, with regards to the estimator of 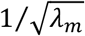, we notice that

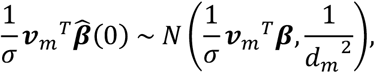

where the terms with larger *m* have larger variance. Thus, we take the average of 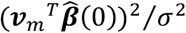, weighted by 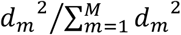, and also subtract the upward bias as,

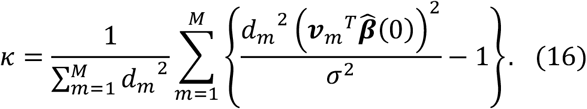

The weighting and subtraction were mentioned in [28], where the subtraction term was dismissed, under the assumption of large effect-size ***β***. Since the effect-size could be small in our application, we keep the subtraction term. The statistic *κ* can be nonpositive, and is unbiased in the sense that

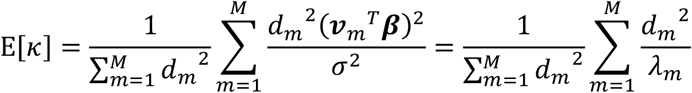

equals the weighted sum of 1/ *λ*_*m*_. Our choice of regularization parameter is

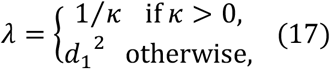

where *d*_1_^2^ is taken instead of positive infinity.

### Implementation of omicwas package

For each omics marker, the parameters ***α, β*** and ***γ*** (denoted in combination by ***θ***) are estimated and tested by nonlinear ridge regression in the following steps. As we assume the magnitude of trait effects ***β*** and ***γ*** to be much smaller than that of basal marker level ***α***, we first fit ***α*** alone for numerical stability.

1. Compute OLS estimator 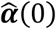 by minimizing formula (6) under ***β*** = ***γ*** = **0**. Apply Wald test.
2. Calculate 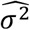 by formula (7). Use it as a substitute for *σ*^2^. The residual degrees of freedom *n* − *p* is the number of samples minus the number of parameters in ***α***.
3. Compute OLS estimators 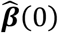 and 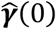 by minimizing formula (6) under 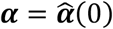. Let 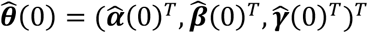.
4. Apply singular value decomposition 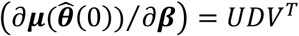.
5. Calculate *κ* and then the regularization parameter *λ* by formulae (16) and (17).
6. Compute ridge estimators 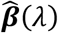 and 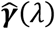 by minimizing formula (10) under 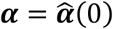. Let 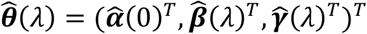.
7. Approximate the variance of ridge estimator, according to formula (12), by

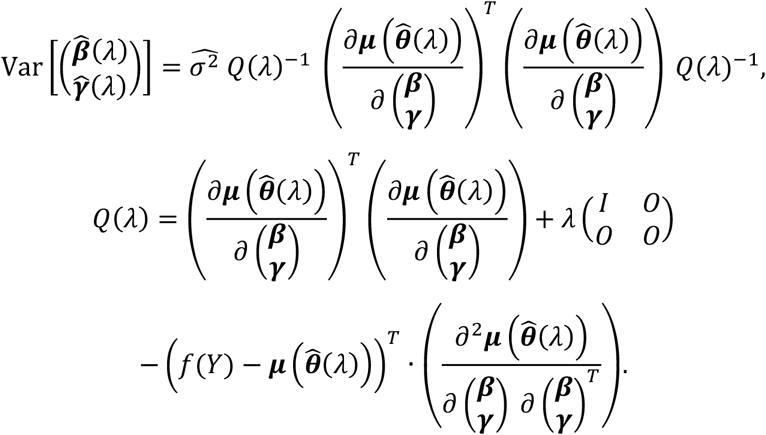
8. Apply the “non-exact” *t*-type test [30]. For the *s*-th coordinate,

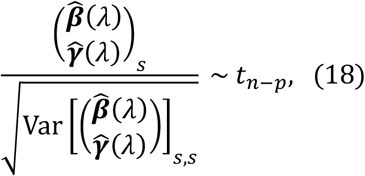

under the null hypothesis 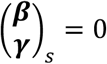.

The formula (18) is the same as a Wald test, but the test differs, because the ridge estimators are not maximum-likelihood estimators. The algorithm was implemented as a package for the R statistical language. We used the NL2SOL algorithm of the PORT library [31] for minimization.

In analyses of quantitative trait locus (QTL), such as methylation QTL (mQTL) and expression QTL (eQTL), an association analysis that takes the genotypes of a single nucleotide polymorphism (SNP) as *X*_*I,k*_ is repeated for many SNPs. In order to speed up the computation, we perform rounds of linear regression. First, the parameters 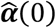 and 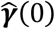 are fit by ordinary linear regression under ***β*** = **0**, which does not depend on *X*_*i,k*_. By taking the residuals, we practically dispense with 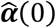 and 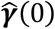 in the remaining steps. Next, for *X*_*i,k*_ of each SNP,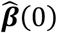 is fit by ordinary linear regression under 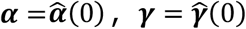. The regularization parameter *λ* is computed according to steps 4 and 5 above. Finally,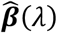is fitted and tested by linear ridge regression under 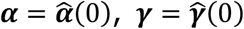.

### Multicollinearity of interaction terms

The regressors for cell-type-specific trait effects in the full model (equation (2)) are the interaction terms *W*_*h,i*_*X*_*i,k*_. To assess multicollinearity, we mathematically derive the correlation coefficient between two interaction terms *W*_*h,i*_*X*_*i,k*_ and *W*_*h*′_ *X*_*i,k*_. In this section, we treat *W*_*h,i*_, *W*_*h*′_ and *X*_*i,k*_ as sampled instances of random variables *W*_*h*_, *W*_*h*′_ and *X*_*k*_, respectively; note that the sample index *i* is omitted. For simplicity, we assume *W*_*h*_ and *W*_*h*′_ are independent of *X*_*k*_. Let E[•], Var[•], Cov[•], Cor[•] and CV[•] denote the expectation, variance, covariance, correlation and coefficient of variation, respectively. Since *X*_*k*_ is centered, E[*W*_*h*_*X*_*k*_] = E[*W*_*h*′_ *X*_*k*_] = 0. The correlation coefficient between interaction terms becomes

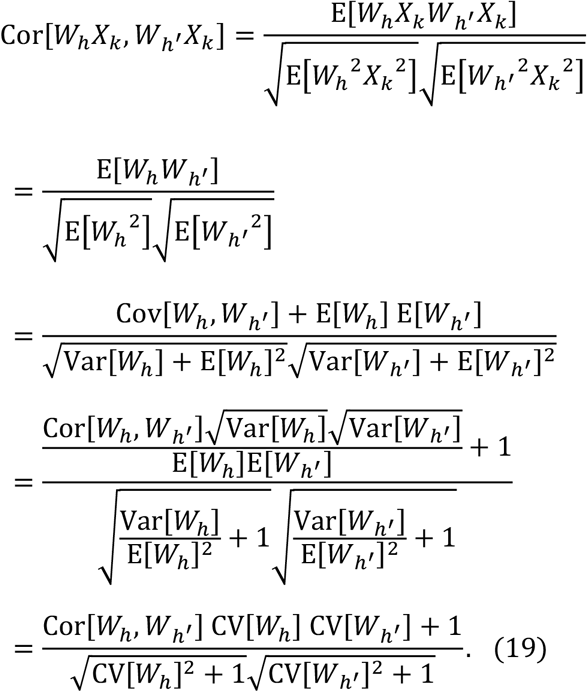

If CV[*W*_*h*_] and CV[*W*_*h*′_] approach zero, the correlation of interaction terms approaches one, irrespective of Cor[*W*_*h*_*M W*_*h*′_].

### EWAS of rheumatoid arthritis and age

EWAS datasets for rheumatoid arthritis were downloaded from the Gene Expression Omnibus. Using the RnBeads package (version 2.2.0) [32] of R, IDAT files of HumanMethylation450 array were preprocessed by removing low quality samples and markers, by normalizing methylation level, and by removing markers on sex chromosomes and outlier samples. The association of methylation level with disease status was tested with adjustment for sex, age, smoking status and experiment batch; the covariates were assumed to have uniform effects across cell types. Alternatively, the association of methylation level with age was tested with adjustment for disease status, sex, smoking status and experiment batch. After quality control, dataset GSE42861 included bulk peripheral blood leukocyte data for 336 cases and 322 controls [22].

The cell type composition of bulk samples was imputed using the Houseman algorithm [33] in the GLINT software (version 1.0.4) [34]. The reference data of GLINT software characterizes seven cell types [35] by 300 CpG sites [36], of which 284 were measured in our data. We used prediction results for the seven cell types (Table 1).

Dataset GSE131989 included sorted CD14^+^ monocyte data for 63 cases and 31 controls [23]. By meta-analysis of GSE131989 and GSE87095 [21], we obtained sorted CD19^+^ B cell data for 108 cases and 95 controls. Under the nominal significance level P < 0.05 (two-sided), the number of CpG sites up- or down-regulated in cases were 20,869 (5%) and 14,911 (3%), respectively, in CD14^+^ monocyte and 28,004 (6%) and 26,582 (6%) in CD19^+^ B cell.

From the Gene Expression Omnibus dataset GSE56047 [25], we obtained sorted CD14^+^ monocyte data for 1200 samples and sorted CD4^+^ T cell data for 214 samples. Under the nominal significance level P < 0.05 (two-sided), the number of CpG sites up- or down-regulated by higher age were 45,283 (10%) and 80,871 (18%), respectively, in CD14^+^ monocyte and 35,822 (8%) and 25,020 (5%) in CD4^+^ T cell.

### Differential gene expression by age

Whole blood RNA-seq data of GTEx v7 was downloaded from the GTEx website [24]. Genes of low quality or on sex chromosomes were removed, expression level was normalized, outlier samples were removed, and 389 samples were retained. The association of read count with age was tested with adjustment for sex.

The cell type composition of bulk samples was imputed using the DeconCell package (version 0.1.0) [10] of R. The reference data of DeconCell characterizes 33 cell types by two to 217 signature genes. Our data measured 39% of the signature genes. We used prediction results for six main cell types (Table 2), for which the prediction performance was 44.4 to 90.9.

From the Gene Expression Omnibus dataset GSE56047 [25], we obtained sorted CD14^+^ monocyte data for 1202 samples and sorted CD4^+^ T cell data for 214 samples. Under the nominal significance level P < 0.05 (two-sided), the number of genes up- or down-regulated by higher age were 2715 (11%) and 3240 (13%), respectively, in CD14^+^ monocyte and 1082 (4%) and 1246 (5%) in CD4^+^ T cell.

### Simulation of cell-type-specific disease association

Bulk tissue sample data for case-control comparison were simulated based on the above-mentioned EWAS dataset GSE42861 and GTEx gene expression dataset. We randomly assigned the case-control status to the samples. Among the omics markers, 2.5% were set to be up-regulated in cases in single cell type, 2.5% were similarly down-regulated, and 95% were unrelated to case-control status. The cell-type-specific effect-size of the differentially expressed markers was fixed within a simulation trial, and was chosen from methylation OR of 1.3, 1.6 or 1.9 for EWAS [20] and fold-change of 1.7, 3.0 or 5.0 for gene expression analysis; the effect-sizes correspond to log(1.3), log(1.6) and log(1.9) or log(1.7), log(3.0) and log(5.0) in normalized scale. If the mean methylation level of a CpG site in cases and controls are *μ*_case_ and *μ*_control_, respectively, the methylation odds become *μ*_case_/ (1 – *μ*_case_) and *μ*_control_/ (1 − *μ*_control_.) The methylation OR represents the case-control contrast of methylation level by the ratio of odds, {*μ*_case_/ (1 – *μ*_case_)}/ {*μ*_control_/ (1 – *μ*_control_)} (see [20]).

For each effect-size, we performed 50 simulation trials. In each simulation trial, we randomly assigned half of the samples as cases (*X*_*i,k*_ = 0.5) and the other half as controls (*X*_*i,k*_ = −0.5). We retained the covariates matrix *C*_*i,l*_ and the cell type composition matrix *W*_*h,i*_ from the original data. From the original bulk expression level matrix *Y*_*i,j*_, 95% of the markers were randomly chosen and retained; these markers had no association with disease because the case-control status was randomized. Cell-type-specific association was introduced into the remaining 5% of markers, such that an equal number of markers were up- or down-regulated in each cell type. For example, in the EWAS dataset, 451,725 × 0.05 × 0.5 ÷ 7 = 1613 CpG sites were up-regulated in neutrophils of cases.

The bulk expression level of a marker *j* with normalized-scale effect-size *β* specific to a cell type *h* was generated as follows. First, the average *μ* and the variance *σ*^2^ of the normalized bulk expression level *f*(*Y*_*i,j*_) in the original data was measured. Next, we generated normalized expression level in each cell type. For cases, the expression level in cell type *h* was randomly sampled from the normal distribution *N*(*μ* + *β, σ*^2^) and the expression level in each of the other cell types was sampled from *N*(*μ, σ*^2^). For controls, the expression level in each cell type was sampled from *N*(*μ, σ*^2^). Finally, for each individual, the expression levels in cell types were converted to the linear scale, multiplied by the cell type composition and added, to obtain the bulk expression level in linear scale.

In the truly disease-associated cell type *h*, we introduced signal *β* and noise *σ*^2^. The signal level was fixed in a simulation trial, for example to methylation OR = 1.3. Since the noise level was taken from real data, the level varied between markers. In the process of obtaining bulk expression the expression of all cell types was mixed, which dilutes the signal. The signal dilution becomes stronger if *h* is a minor cell type. The mixing process adds noise from other cell types, which becomes stronger if *h* is a minor cell type. Consequently, minor cell types tend to manifest weaker association in bulk tissue. We empirically measured the strength of association by the coefficient of determination, *R*^2^, for the marginal model. The coefficient of determination is defined as the proportion of variance explained by the model, and *R*^2^/ (1 − *R*^2^) equals the signal-to-noise ratio. Under methylation OR of 1.3, 1.6, 1.9 for EWAS simulation, the median *R*^2^ was 0.322, 0.589, 0.712 for neutrophils, 0.010, 0.033, 0.057 for NK cells, and 0.001, 0.003, 0.005 for eosinophils. Under fold-change of 1.7, 3.0 or 5.0 for gene expression simulation, the median *R*^2^ was 0.135, 0.331, 0.434 for granulocytes, 0.007, 0.026, 0.049 for CD8^+^ T cells, and 0.001, 0.003, 0.007 for B cells.

For gene expression, we also simulated a scenario where cell-type-specific disease effect occurs in cell type marker genes. The simulation procedure is same as above except that the expression level was set to zero (in linear scale) in all cell types other than the target cell type *h*, for both cases and controls.

### Evaluation of statistical methods

Cell-type-specific effects of traits was statistically tested by using bulk tissue data as input. We applied the omicwas package with the normalizing function *f* = log, logit, identity without ridge regularization (omicwas.log, omicwas.logit, omicwas.identity) or under ridge regression (omicwas.log.ridge, omicwas.logit.ridge, omicwas.identity.ridge). The omicwas package was used also for conventional linear regression under the full and marginal models. We also developed a hybrid of marginal and full models (Marginal.Full005): if the effect direction agreed in two models and if P < 0.05 in the full model, we adopted the Z-score of the marginal model; otherwise, the Z-score was set to zero.

Among previous methods, we evaluated those that accept cell type composition as input and compute test statistics for cell-type-specific association. For DNA methylation data, we applied TOAST (version 1.2.0) [9], CellDMC (version 2.0.2) [13] and TCA (version 1.0.0) [14]. For gene expression data, we applied TOAST and csSAM (version 1.4) [5]. For csSAM, we either fitted all cell types together or one cell type at a time, and denoted the results as csSAM.lm and csSAM.monovariate, respectively. The csSAM method is applicable to binomial traits but not to quantitative traits.

For simulated data of EWAS dataset GSE42861, we adopted the significance level P < 2.4 × 10^−7^, which accounts for the correlation among the probes on HumanMethylation450 array [37]. For the GTEx gene expression dataset, multiple testing was controlled by the Benjamini-Hochberg procedure with the false discovery rate <5% in each cell type [38]. The performance of an algorithm for the simulated data was assessed by sensitivity, specificity and precision. The performance measures were obtained from each simulation trial. For a target cell type *h*, we counted the four possible outcomes, true positives (TP_*h*_), true negatives (TN_*h*_), false positives (FP_*h*_) and false negatives (FN_*h*_). The sum TP_*h*_ + TN_*h*_ + FP_*h*_ + FN_*h*_ equals the total number of omics markers (which was 451,725 CpG sites for DNA methylation and 14,038 genes for gene expression). For 5% of the markers, one randomly selected cell type *h*^∗^ was set to be truly associated with disease status at data generation. The remaining 95% of the markers were null cases with no truly associated cell types. The outcome counts can be subtotaled according to the truly associated cell type, which is denoted in superscript,

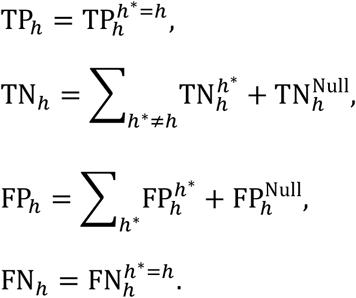

Remark that FP_*h*_ can occur when in cell type *h* a marker is truly up-regulated in disease cases but an algorithm predicts the marker to be down-regulated in *h*. The performance measures can be represented as

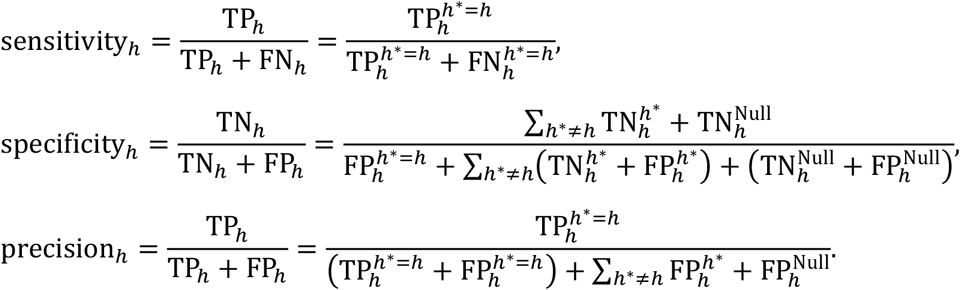

Whereas sensitivity is obtained solely from markers that are truly associated in the target cell type *h*, the specificity and precision are obtained by aggregating with the markers associated in other cell types and the null markers.

For the association with rheumatoid arthritis and age, “true” association was determined from the measurements in physically sorted blood cells, under the nominal significance level P < 0.05 (two-sided). In the same manner as the simulation analysis, we assessed the sensitivity, specificity and precision.

## Supplementary information

**Additional file 1: Table S1**. Blood cell type proportion in Tsimane Amerindians, Caucasians and Hispanics.

**Additional file 2: Fig. S1**. Sensitivity for detecting cell-type-specific association in simulated data for gene expression of marker genes.

**Additional file 3: Fig. S2**. Specificity for detecting cell-type-specific association in simulated data for gene expression of marker genes.

**Additional file 4: Fig. S3**. Precision (positive predictive value) for detecting cell-type-specific association in simulated data for gene expression of marker genes.

**Additional file 5: Supplementary note**. Asymptotic distribution of ridge estimator.

## Abbreviations

CV: coefficient of variation
eQTL: expression QTL
EWAS: epigenome-wide association study
mQTL: methylation QTL
MSE: mean squared error
OLS: ordinary least squares
OR: odds ratio
QTL: quantitative trait locus
SNP: single nucleotide polymorphism

## Declarations

### Ethics approval and consent to participate

Not applicable.

### Consent for publication

Not applicable.

### Availability of data and materials

The datasets generated and analyzed during the current study are available in the figshare repository, https://dx.doi.org/10.6084/m9.figshare.10718282

## Author’s contributions

FT developed the methodology, wrote the software, implemented the study, and wrote the manuscript. NK revised the manuscript. All authors read and approved the final manuscript.

## Acknowledgements

Not applicable.

## References

1. Teschendorff AE, Zheng SC. Cell-type deconvolution in epigenome-wide association studies: a review and recommendations. Epigenomics. 2017;9:757–68.

2. Sturm G, Finotello F, Petitprez F, Zhang JD, Baumbach J, Fridman WH, et al. Comprehensive evaluation of transcriptome-based cell-type quantification methods for immuno-oncology. Bioinformatics. 2019;35:i436–45.

3. Ghosh D. Mixture models for assessing differential expression in complex tissues using microarray data. Bioinformatics. 2004;20:1663–9.

4. Stuart RO, Wachsman W, Berry CC, Wang-Rodriguez J, Wasserman L, Klacansky I, et al. In silico dissection of cell-type-associated patterns of gene expression in prostate cancer. Proc Natl Acad Sci USA. National Academy of Sciences; 2004;101:615–20.

5. Shen-Orr SS, Tibshirani R, Khatri P, Bodian DL, Staedtler F, Perry NM, et al. Cell type–specific gene expression differences in complex tissues. Nat Meth. Nature Publishing Group; 2010;7:287–9.

6. Erkkilä T, Lehmusvaara S, Ruusuvuori P, Visakorpi T, Shmulevich I, Lähdesmäki H. Probabilistic analysis of gene expression measurements from heterogeneous tissues. Bioinformatics. 2010;26:2571–7.

7. Kuhn A, Thu D, Waldvogel HJ, Faull RLM, Luthi-Carter R. Population-specific expression analysis (PSEA) reveals molecular changes in diseased brain. Nat Meth. Nature Publishing Group; 2011;8:945–7.

8. Westra H-J, Arends D, Esko T, Peters MJ, Schurmann C, Schramm K, et al. Cell Specific eQTL Analysis without Sorting Cells. Pastinen T, editor. PLoS Genet. Public Library of Science; 2015;11:e1005223–17.

9. Li Z, Wu Z, Jin P, Wu H. Dissecting differential signals in high-throughput data from complex tissues. Hancock J, editor. Bioinformatics. 2019;35:3898–905.

10. Aguirre-Gamboa R, de Klein N, di Tommaso J, Claringbould A, van der Wijst MG, de Vries D, et al. Deconvolution of bulk blood eQTL effects into immune cell subpopulations. BMC Bioinformatics. 2020;21:243.

11. Montaño CM, Irizarry RA, Kaufmann WE, Talbot K, Gur RE, Feinberg AP, et al. Measuring cell-type specific differential methylation in human brain tissue. Genome Biol. BioMed Central; 2013;14:R94–9.

12. White N, Benton M, Kennedy D, Fox A, Griffiths L, Lea R, et al. Accounting for cell lineage and sex effects in the identification of cell-specific DNA methylation using a Bayesian model selection algorithm. Sawalha AH, editor. PLoS ONE. 2017;12:e0182455–18.

13. Zheng SC, Breeze CE, Beck S, Teschendorff AE. Identification of differentially methylated cell types in epigenome-wide association studies. Nat Meth. Nature Publishing Group; 2018;15:1059–66.

14. Rahmani E, Schweiger R, Rhead B, Criswell LA, Barcellos LF, Eskin E, et al. Cell-type-specific resolution epigenetics without the need for cell sorting or single-cell biology. Nature Communications. Nature Publishing Group; 2019;10:3417–11.

15. Cobos FA, Vandesompele J, Mestdagh P. Computational deconvolution of transcriptomics data from mixed cell populations. Bioinformatics. 2018;34:1969–79.

16. Hoyle DC, Rattray M, Jupp R, Brass A. Making sense of microarray data distributions. Bioinformatics. 2002;18:576–84.

17. Du P, Zhang X, Huang C-C, Jafari N, Kibbe WA, Hou L, et al. Comparison of Beta-value and M-value methods for quantifying methylation levels by microarray analysis. BMC Bioinformatics. BioMed Central; 2010;11:1–9.

18. Zhuang J, Widschwendter M, Teschendorff AE. A comparison of feature selection and classification methods in DNA methylation studies using the Illumina Infinium platform. BMC Bioinformatics. BioMed Central; 2012;13:1–14.

19. Aiken LS, West SG. Multiple Regression: Testing and Interpreting Interactions. Sage Publications; 1991.

20. Rakyan VK, Down TA, Balding DJ, Beck S. Epigenome-wide association studies for common human diseases. Nat Rev Genet. 2011;12:529–41.

21. Julià A, Absher D, López-Lasanta M, Palau N, Pluma A, Waite Jones L, et al. Epigenome-wide association study of rheumatoid arthritis identifies differentially methylated loci in B cells. Hum Mol Genet. 2017;26:2803–11.

22. Liu Y, Aryee MJ, Padyukov L, Fallin MD, Hesselberg E, Runarsson A, et al. Epigenome-wide association data implicate DNA methylation as an intermediary of genetic risk in rheumatoid arthritis. Nat Biotechnol. Nature Publishing Group; 2013;31:142–7.

23. Rhead B, Holingue C, Cole M, Shao X, Quach HL, Quach D, et al. Rheumatoid Arthritis Naive T Cells Share Hypermethylation Sites With Synoviocytes. Arthritis & Rheumatology. John Wiley & Sons, Ltd; 2017;69:550–9.

24. GTEx Consortium. Genetic effects on gene expression across human tissues. Nature. Nature Publishing Group; 2017;550:204–13.

25. Reynolds LM, Taylor JR, Ding J, Lohman K, Johnson C, Siscovick D, et al. Age-related variations in the methylome associated with gene expression in human monocytes and T cells. Nature Communications. 2014;5:5366.

26. Riazoshams H, Midi H, Ghilagaber G. Robust Nonlinear Regression: with Applications using R. John Wiley & Sons; 2019.

27. Lim C. Robust ridge regression estimators for nonlinear models with applications to high throughput screening assay data. Statist. Med. 2014;34:1185–98.

28. Lawless JF, Wang P. A simulation study of ridge and other regression estimators. Communications in Statistics -Theory and Methods. 1976;5:307–23.

29. Hoerl AE, Kannard RW, Baldwin KF. Ridge regression: some simulations. Communications in Statistics - Theory and Methods. 1975;4:105–23.

30. Halawa AM, Bassiouni El MY. Tests of regression coefficients under ridge regression models. Journal of Statistical Computation and Simulation. 2000;65:341–56.

31. Dennis JE, Gay DM, Welsch RE. An adaptive nonlinear least-squares algorithm. ACM Transactions on Mathematical Software. 1981;7:348–68.

32. Müller F, Scherer M, Assenov Y, Lutsik P, Walter J, Lengauer T, et al. RnBeads 2.0: comprehensive analysis of DNA methylation data. Genome Biol. BioMed Central; 2019;20:55–12.

33. Houseman EA, Accomando WP, Koestler DC, Christensen BC, Marsit CJ, Nelson HH, et al. DNA methylation arrays as surrogate measures of cell mixture distribution. BMC Bioinformatics. 2012;13:86.

34. Rahmani E, Yedidim R, Shenhav L, Schweiger R, Weissbrod O, Zaitlen N, et al. GLINT: a user-friendly toolset for the analysis of high-throughput DNA-methylation array data. Hancock JM, editor. Bioinformatics. 2017;33:1870–2.

35. Reinius LE, Acevedo N, Joerink M, Pershagen G, Dahlén S-E, Greco D, et al. Differential DNA Methylation in Purified Human Blood Cells: Implications for Cell Lineage and Studies on Disease Susceptibility. Ting AH, editor. PLoS ONE. Public Library of Science; 2012;7:e41361–13.

36. Koestler D. Improving Cell Mixture Deconvolution by Identifying Optimal DNA methylation Libraries (IDOL). BMC Bioinformatics. BMC Bioinformatics; 2016;:1–21.

37. Saffari A, Silver MJ, Zavattari P, Moi L, Columbano A, Meaburn EL, et al. Estimation of a significance threshold for epigenome-wide association studies. Genet Epidemiol. John Wiley & Sons, Ltd; 2017;42:20–33.

38. Anders S, McCarthy DJ, Chen Y, Okoniewski M, Smyth GK, Huber W, et al. Count-based differential expression analysis of RNA sequencing data using R and Bioconductor. Nat Protoc. 2013;8:1765–86.

