## Additional file 2: Fig. S1. for "Nonlinear ridge regression improves cell-type-specific differential expression analysis"

Figure S1

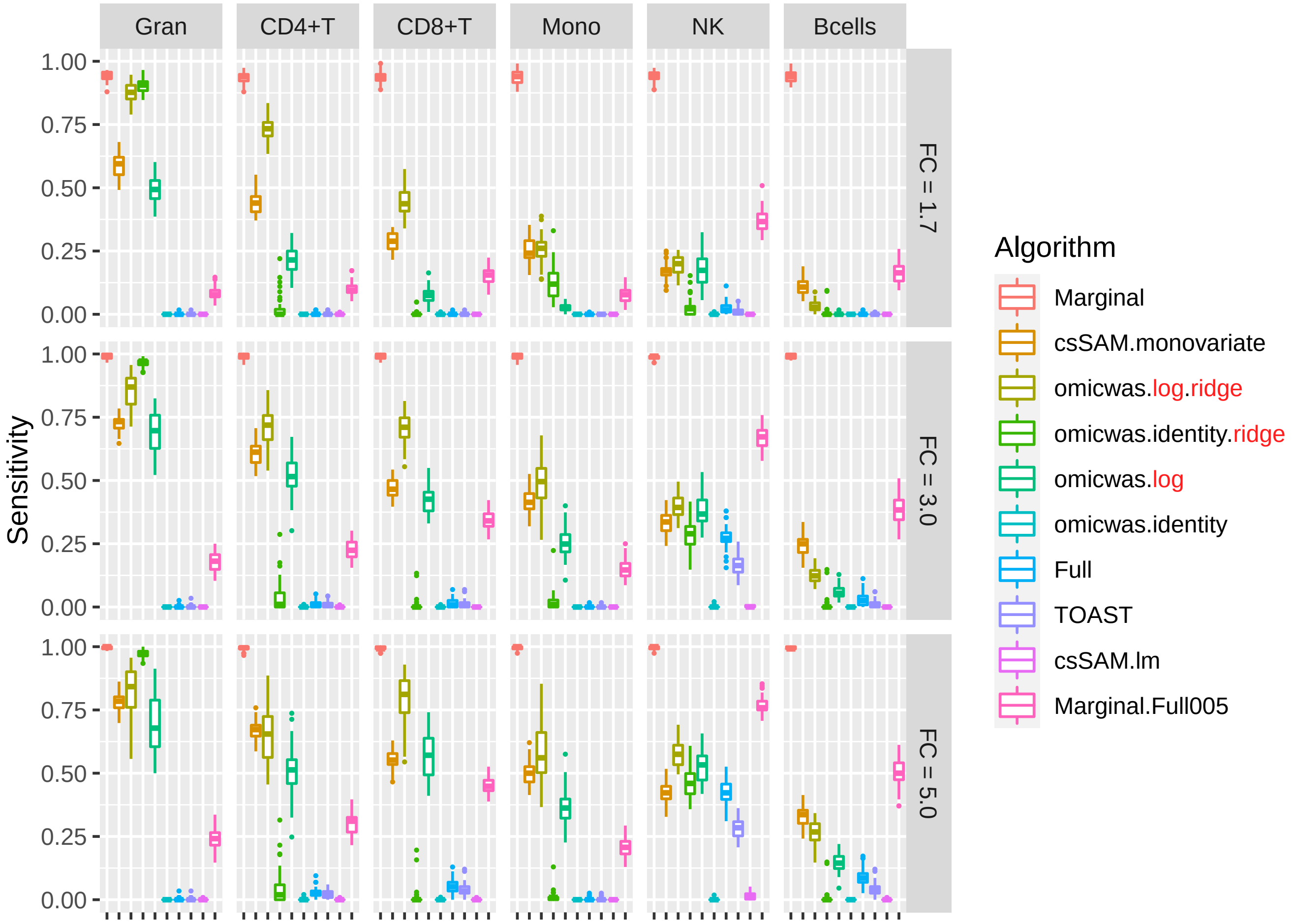


Sensitivity for detecting cell-type-specific association in simulated data for gene expression of marker genes. Panels are aligned in rows according to the simulation settings with the gene expression fold change of 1.7, 3.0 or 5.0. In each row, panels for different cell types are aligned in decreasing order of proportion. The vertical axis indicates sensitivity. In each panel, results from different algorithms are aligned horizontally in different colors. Results from 50 simulation trials are summarized in a box plot. The middle bar of the box plot indicates the median, and the lower and upper hinges correspond to the first and third quartiles. The whiskers extend to the value no further than 1.5 x inter-quartile range from the hinges. FC, fold change; Gran, granulocytes; Mono, monocytes.
