## Additional file 3: Fig. S2. for "Nonlinear ridge regression improves cell-type-specific differential expression analysis"

Figure S2

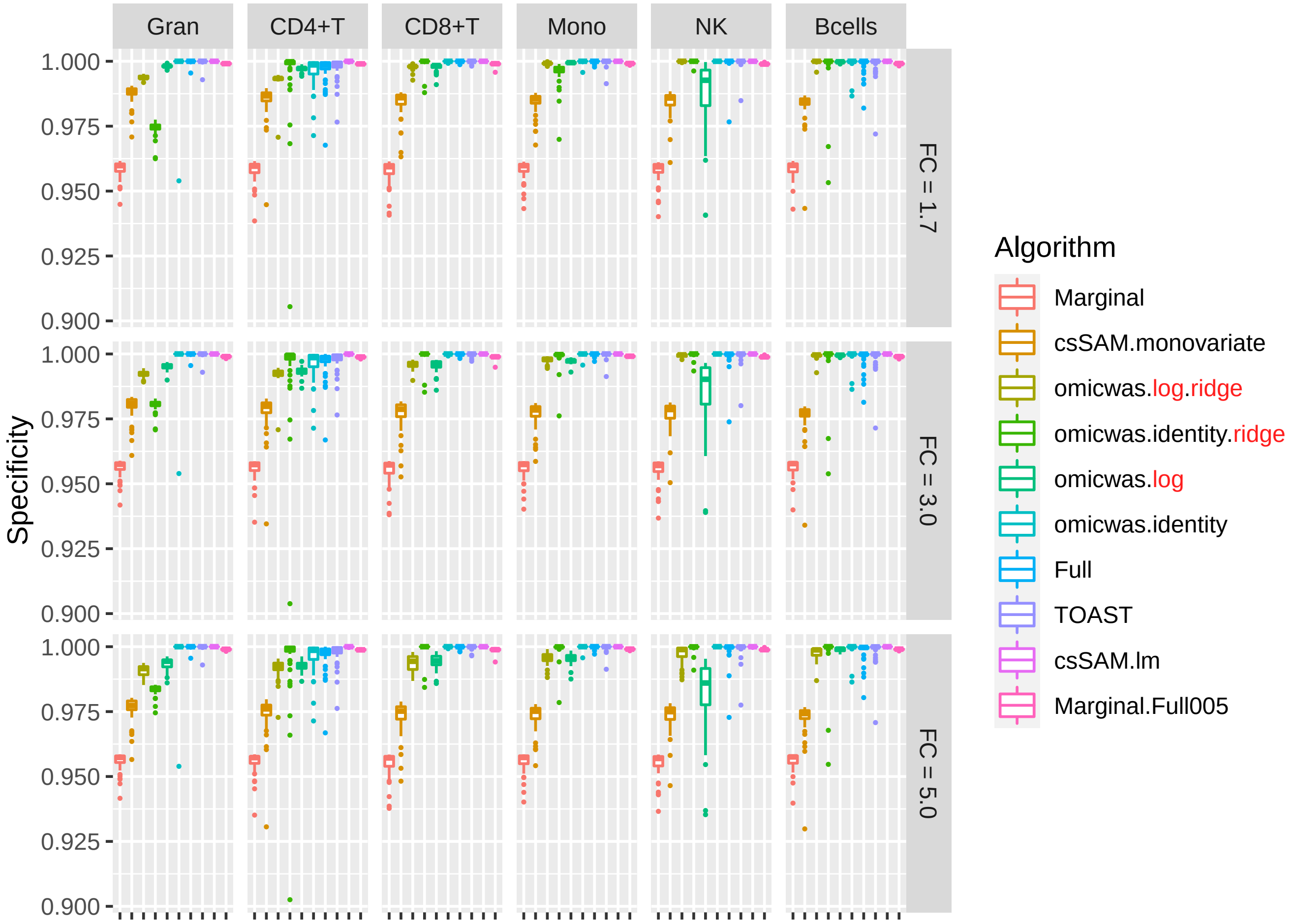


Specificity for detecting cell-type-specific association in simulated data for gene expression of marker genes. The figure format is same as Fig. S1.
