## Additional file 4: Fig. S3. for "Nonlinear ridge regression improves cell-type-specific differential expression analysis"

Figure S3

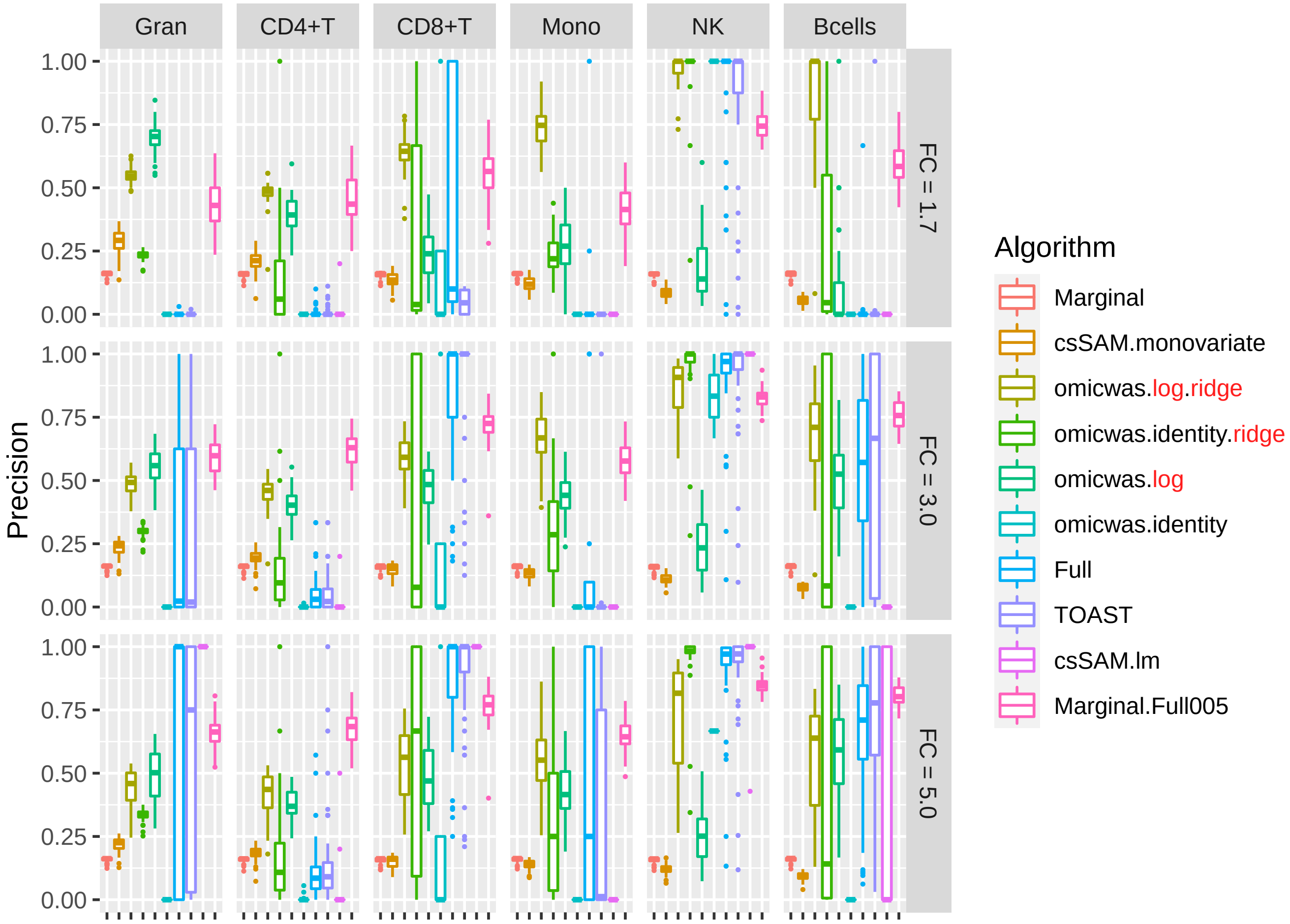


Precision (positive predictive value) for detecting cell-type-specific association in simulated data for gene expression of marker genes. The figure format is same as Fig. S1.
