## Additional file 5: Supplementary note. for "Nonlinear ridge regression improves cell-type-specific differential expression analysis"

### Asymptotic distribution of ridge estimator

We here show that the ridge estimator $\widehat{\boldsymbol{\theta}}(\lambda)$ is asymptotically normally distributed with the mean and variance in equations (11) and (12). Since the ridge estimator minimizes formula (10), its partial derivatives with respect to single parameters equal zero:

$$\left(\frac{\partial \boldsymbol{\mu}\left(\widehat{\boldsymbol{\theta}}(\lambda)\right)}{\partial \boldsymbol{\theta}}\right)^T \left(f(Y) - \boldsymbol{\mu}\left(\widehat{\boldsymbol{\theta}}(\lambda)\right)\right) - \lambda \begin{pmatrix} O & O & O \\ O & I & O \\ O & O & O \end{pmatrix} \widehat{\boldsymbol{\theta}}(\lambda) = \mathbf{0}. \quad \text{(A1)}$$

The Taylor series of $\boldsymbol{\mu}$ and its Jacobian with regards to the variable $\widehat{\boldsymbol{\theta}}(\lambda)$ centered at the true parameter value $\boldsymbol{\theta}$ become

$$\boldsymbol{\mu}\left(\widehat{\boldsymbol{\theta}}(\lambda)\right) \approx \boldsymbol{\mu}(\boldsymbol{\theta}) + \left(\frac{\partial \boldsymbol{\mu}(\boldsymbol{\theta})}{\partial \boldsymbol{\theta}}\right)\left(\widehat{\boldsymbol{\theta}}(\lambda) - \boldsymbol{\theta}\right),$$

$$\left(\frac{\partial \boldsymbol{\mu}\left(\widehat{\boldsymbol{\theta}}(\lambda)\right)}{\partial \boldsymbol{\theta}}\right) \approx \left(\frac{\partial \boldsymbol{\mu}(\boldsymbol{\theta})}{\partial \boldsymbol{\theta}}\right) + \left(\frac{\partial^2 \boldsymbol{\mu}(\boldsymbol{\theta})}{\partial \boldsymbol{\theta} \partial \boldsymbol{\theta}^T}\right) \cdot \left(\widehat{\boldsymbol{\theta}}(\lambda) - \boldsymbol{\theta}\right),$$

where the dot product of $(\partial^2 \boldsymbol{\mu}(\boldsymbol{\theta}) / \partial \boldsymbol{\theta} \partial \boldsymbol{\theta}^T)$ and the next term is taken by multiplying for each parameter and then summing up over parameters. We neglect the second or higher order terms of $\widehat{\boldsymbol{\theta}}(\lambda) - \boldsymbol{\theta}$. By plugging into equation (A1), we obtain

$$\left(\frac{\partial \boldsymbol{\mu}(\boldsymbol{\theta})}{\partial \boldsymbol{\theta}}\right)^T \left(f(Y) - \boldsymbol{\mu}(\boldsymbol{\theta})\right) - \left\{\left(\frac{\partial \boldsymbol{\mu}(\boldsymbol{\theta})}{\partial \boldsymbol{\theta}}\right)^T \left(\frac{\partial \boldsymbol{\mu}(\boldsymbol{\theta})}{\partial \boldsymbol{\theta}}\right) - \left(f(Y) - \boldsymbol{\mu}(\boldsymbol{\theta})\right)^T \cdot \left(\frac{\partial^2 \boldsymbol{\mu}(\boldsymbol{\theta})}{\partial \boldsymbol{\theta} \partial \boldsymbol{\theta}^T}\right)\right\}\left(\widehat{\boldsymbol{\theta}}(\lambda) - \boldsymbol{\theta}\right)$$

$$-\lambda \begin{pmatrix} O & O & O \\ O & I & O \\ O & O & O \end{pmatrix} \widehat{\boldsymbol{\theta}}(\lambda) = \mathbf{0}.$$

Thus,

$$\widehat{\boldsymbol{\theta}}(\lambda) = Q(\lambda)^{-1}\, Q(0)\, \boldsymbol{\theta} + Q(\lambda)^{-1} \left(\frac{\partial \boldsymbol{\mu}(\boldsymbol{\theta})}{\partial \boldsymbol{\theta}}\right)^T \left(f(Y) - \boldsymbol{\mu}(\boldsymbol{\theta})\right)$$

$$= Q(\lambda)^{-1}\, Q(0)\, \boldsymbol{\theta} + Q(\lambda)^{-1} \left(\frac{\partial \boldsymbol{\mu}(\boldsymbol{\theta})}{\partial \boldsymbol{\theta}}\right)^T \boldsymbol{\varepsilon},$$

where $\boldsymbol{\varepsilon} \sim N(\mathbf{0}, \sigma^2 I)$.
